# Activation of the protective arm of renin-angiotensin system enhances mitochondrial turnover improving respiration and decreasing integrated stress response in a human Complex III deficiency model

**DOI:** 10.64898/2026.03.20.711686

**Authors:** Lucía Fernández-del-Río, Andrea Eastes, David Rincón Fernández Pacheco, Nikole Scillitani, Jasmine Garza, Matthew Dugan, Matheus Pinto de Oliveira, Pradnya Kadam, Iman Gauhar, Karel Erion, Kathleen Rodgers, Kevin Gaffney, Amy Wang, Marc Liesa, Cristiane Benincá, Orian Shaul Shirihai

## Abstract

Primary mitochondrial diseases are clinically and genetically heterogeneous disorders, commonly caused by defects in the oxidative phosphorylation system. This heterogeneity presents major challenges for therapeutic development; however, a shared hallmark across these diseases is the accumulation of dysfunctional mitochondria. Enhancing mitochondrial turnover, by activating the selective degradation of dysfunctional mitochondria via mitophagy, concurrently with the activation of mitochondrial biogenesis, could represent a shared therapeutic strategy for mitochondrial diseases. Here, we describe a novel mitophagy inducer, CAP-1902. CAP-1902 is a new agonist of the MAS G-Protein Coupled Receptor (MasR). In fibroblasts from patients carrying a *BCS1L* mutation that impairs complex III (CIII) assembly, CAP-1902 increased mitochondrial turnover by promoting both mitophagy and biogenesis. Specifically, MasR activation triggered the AMPK/ULK1/FUNDC1 mitophagy pathway. Knockdown of FUNDC1 blocked mitophagy but not AMPK activation, confirming pathway specificity. Additionally, a decrease in the occurrence of depolarized mitochondria with treatment indicated the selective targeting of accumulated damaged mitochondria in the disease context. MasR activation by CAP-1902 also stimulated the nuclear translocation of PGC-1α, promoting increased expression of transcripts associated with mitochondrial biogenesis, respiratory chain components, and mitochondrial translation. Remarkably, CAP-1902 was ultimately able to restore key defects in CIII-deficient fibroblasts by rescuing bioenergetics and correcting both the aberrant lysosomal distribution and the elevated integrated stress response markers, which is consistent with a shift toward a healthier mitochondrial population. In summary, we describe the first potential GPCR-mediated treatment of mitochondrial diseases and demonstrate that MasR activation by CAP-1902 induces mitochondrial turnover and improves mitochondrial function.

## INTRODUCTION

Mitochondrial diseases affect approximately 1 in every 4,300 individuals in the United States. Despite this prevalence, only recently has the U.S. Food and Drug Administration approved therapies for specific ultra-rare conditions, including Barth syndrome and Thymidine kinase 2 deficiency. These approvals represent a major milestone in translating decades of mitochondrial research into clinical therapies; however, most mitochondrial disorders still lack disease-specific treatments. The accumulation of dysfunctional mitochondria is a hallmark of primary mitochondrial diseases and is associated with the upregulation of mitophagy-related pathways, often causing mitophagy flux impairment [1]. Simply promoting mitophagy without identifying and correcting the specific point of dysfunction can exacerbate the impairment in mitochondrial clearance. Moreover, enhancing the degradation of damaged mitochondria without simultaneously supporting mitochondrial biogenesis may lead to a net depletion of mitochondria, further compromising cellular bioenergetics. Mitochondrial turnover emerges as a promising target in the therapeutic approach to mitochondrial diseases, serving as a key strategy to uphold mitochondrial homeostasis. The elimination of dysfunctional organelles, coupled with the generation of more efficient ATP producers, stands out as a crucial intervention in these OXPHOS-deficient conditions [2–6]. Indeed, modelling of the mitochondrial life cycle predicts that an elevated number of iterations of this cycle will improve the quality of the overall population [7]. Thus, it is expected that increased turnover can generate and retain a population of mitochondria that successfully adapts and at least mitigates the stress response imposed by genetic disturbances in the OXPHOS machinery. The more cycles that occurred, the higher the chance of preserving a healthier population.

This study provides evidence that the Renin-Angiotensin System (RAS) is a regulator of mitochondrial turnover and holds therapeutic promise for mitochondrial diseases. RAS consists of two opposing arms: the classical pathway and the protective arm. In the classical arm, the primary receptor is the Angiotensin type 1 receptor (AT1R) and the pathway is regulated by angiotensin-converting enzyme [8] and the active effector Angiotensin II (Ang II). In the protective arm, the main receptors are Ang type 2 receptor (AT1R) and Mas receptor (MasR), and the pathway is regulated by angiotensin-converting enzyme ACE2 and the active effector Ang-(1-7). While classical RAS inhibitors remain central to treating cardiovascular and renal disease, recent mechanistic and clinical studies, accelerated by insights from the COVID-19 pandemic, have refocused attention on the protective arm of RAS [9, 10].

Specifically, Mas receptor (MasR) activators, most notably the small-molecule candidate BIO101 (Sarconeos), are advancing through clinical development for age-related sarcopenia, with emerging evidence indicating that MasR signaling can regulate mitochondrial function and bioenergetics resilience [11, 12]. While Ang II has been shown to impair autophagy and mitochondrial function [13], the impact of the protective arm of RAS on mitochondrial function remains poorly explored. The impact of MasR activation on mitochondrial function was mainly studied by focusing on its role in counterbalancing oxidative stress by antagonizing the effect of Ang II/AT1R signaling axis [14]. MasR activation was shown to increase mitochondrial mass and respiration [15] and decrease mitochondrial superoxide [16], effects that could be mediated by the induction of mitochondrial turnover.

Under normal physiological conditions, mitochondrial turnover is controlled by the balance of two interlinked processes, mitophagy and biogenesis. Mitophagy begins with the recognition and engulfment of dysfunctional mitochondria by autophagosomes, followed by their delivery to and degradation within lysosomes [17]. Targeting of mitochondria for degradation can be either initiated by ubiquitin-dependent pathways or receptor-mediated mechanisms involving proteins like PINK, PARKIN, BNIP3, NIX, or FUNDC1. In parallel, mitochondrial biogenesis relies on coordinated transcription of nuclear and mitochondrial genomes driven by the coactivator PGC1α. Energy sensors like AMPK and SIRT1 govern PGC1α activity [18]; activated AMPK not only enhances biogenesis but also triggers mitophagy by phosphorylating ULK1 [19, 20], which in turn phosphorylates FUNDC1 to initiate autophagosome formation around the targeted mitochondria. AMPK is one of the master regulators of cellular and mitochondrial homeostasis, and its functions are linked with the regulation of mitochondrial turnover [21]. Several stimuli can activate AMPK, including the activation of MasR by Ang-(1-7), its endogenous ligand [22].

Here, we describe a novel role for MasR in mitochondrial turnover by activating the receptor with a new agonist (CAP-1902) in a cellular model of human CIII-deficiency. Activation of MasR by CAP-1902 induces selective mitophagy via the ULK1/FUNDC1 pathway and mitochondrial biogenesis through PGC1α activation. The combined removal of depolarized mitochondria and generation of new, more efficient organelles leads to decreased cellular stress and partial restoration of the deficient respiration observed in this model of primary mitochondrial disease. This study establishes a promising strategy for treating mitochondrial disease and demonstrates the benefits of inducing mitochondrial turnover at both the organelle and cellular levels.

## RESULTS

### Activation of MasR induces mitochondrial turnover by selectively targeting depolarized mitochondria

The role of MasR on mitochondrial function has been poorly explored, focusing mainly on its role counterbalancing oxidative stress by antagonizing the effect of angiotensin II (Ang II)/AT1R signaling axis [14]. MasR activation by Ang-(1-7) was shown to increase respiration [15] and decrease mitochondrial superoxide [16], effects that could be mediated by the induction of mitochondrial turnover. To test this hypothesis, we treated fibroblasts derived from control (Ctrl) subjects and OXPHOS-deficient patients with Ang-(1-7) and analyzed mitochondrial summed area and morphology (Figure 1A-B and S1A-C). The OXPHOS deficient fibroblasts are derived from patients with mutations causing complex V (MT-ATP6^L156R^) and complex III (P1:BCS1L^R73C/F368I^ and P2:BCS1L^R183C/R184C^) deficiencies. After 16 h of incubation with Ang-(1–7), mitochondrial morphology and summed area were assessed by confocal microscopy in fibroblasts fixed and stained with an anti-GRP75 antibody to label the mitochondrial matrix (Figures 1A-B and S1A-C). GRP75 labeling was used to enhance spatial resolution and enable accurate segmentation of the mitochondrial network. While mitochondrial morphology was not changed (Figure S1A-C), a decrease in mitochondrial matrix summed area (Figure 1A-B) was observed with Ang-(1-7) treatment in Ctrl cells and the CV-deficient due to *MT-ATP6* mutation (ATP6^L156R^). Interestingly, the ATP6^L156R^ cells showed an increase in summed matrix area compared with Ctrl fibroblasts. No changes were observed following treatment of CIII-deficient (P1:BCS1L^R73C/F368I^ and P2:BCS1L^R183C/R184C^) cells, which already exhibited reduced mitochondrial summed matrix area compared with Ctrl cells. Further validation using an anti-TOMM20 antibody to label the mitochondrial outer membrane revealed no differences in mitochondrial outer membrane summed area in Ctrl cells treated with Ang-(1-7), whereas a slight decrease was observed for the CV-deficient cells (ATP6^L156R^) (Figure S1D-E) that was consistent with the decrease in matrix summed area. These findings can reflect differences in protein abundance in the matrix and the outer membrane, differences in spatial resolution during acquisition, or differential changes in mitochondrial outer membrane area versus matrix area. To clarify these aspects, an anti-dsDNA antibody was used to label mtDNA to determine which of the changes could be consistent with a coordinated decrease in mitochondrial mass. To clarify these aspects, an anti-dsDNA antibody was used to label mtDNA to determine which of the changes could be consistent with a coordinated decrease in mitochondrial mass. CV-deficient cells treated with Ang-(1-7) showed a decrease in mtDNA-stained area, indicating a potential induction of mitochondria clearance and thus decreased mitochondrial mass in treated CV-deficient cells and not in treated control cells (Figure S1F-G). While an increase in mitochondrial mass typically indicates mitochondrial biogenesis, a decrease in mitochondrial mass and mtDNA area suggest enhanced mitophagy flux.

**Figure 1.**
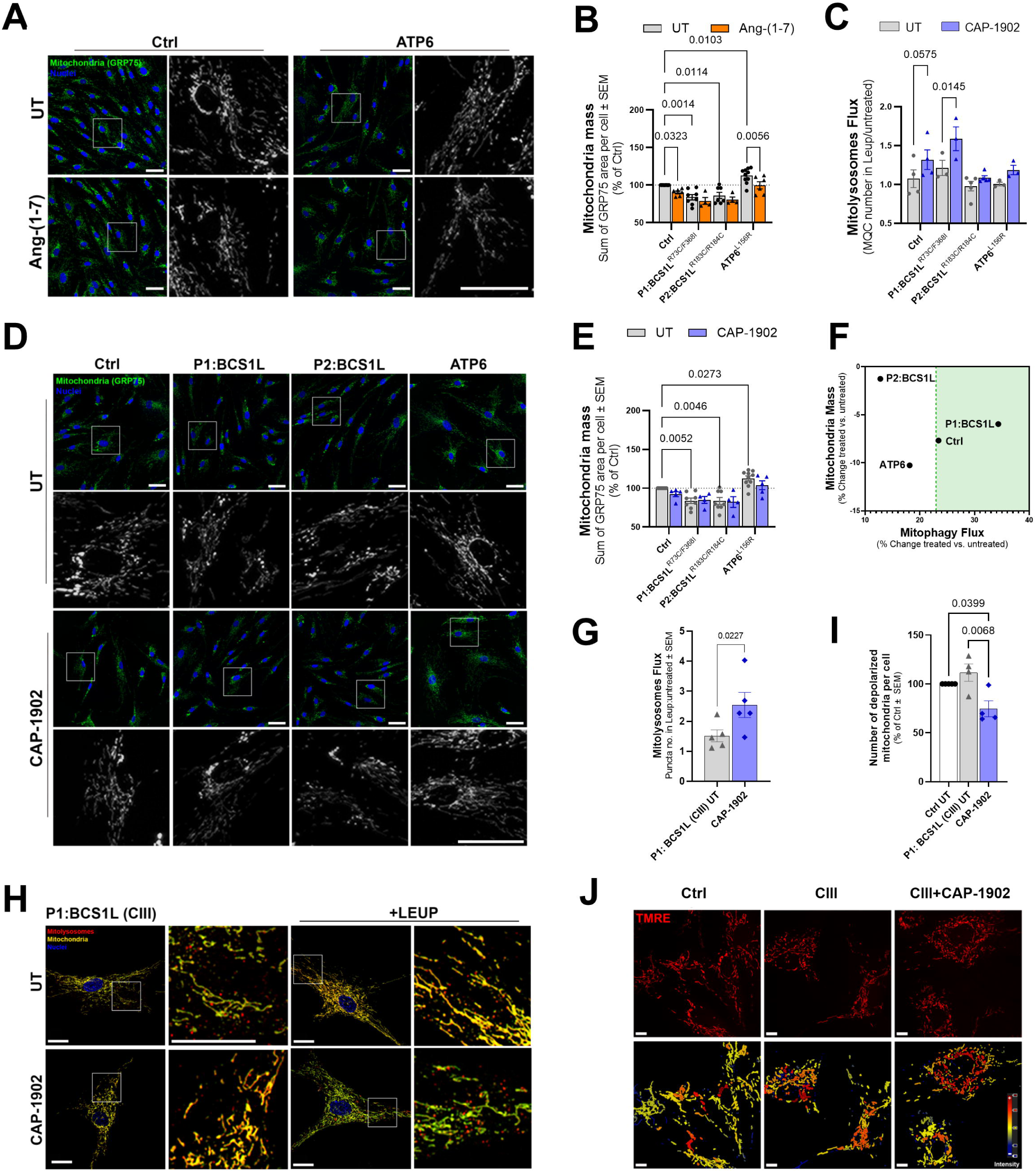
MasR agonists increase mitochondrial turnover in mitochondrial disease patient-derived fibroblasts. (**A**) Representative high-throughput confocal micrographs of Control subjects (Ctrl) and *MT-ATP6* patient mutant (ATP6) fibroblasts labeled with GRP75 antibody (mitochondria - green), and DAPI (nuclei – blue). Cells were treated with 500 nM Ang-(1-7) for 16 h. Scale bars: 50 μm. **(B)** Quantification of mitochondrial mass as sum of area per cell from GRP75 images. Statistical analysis was performed by two-way ANOVA. **(C)** Number of mitolysosomes per cell analyzed in mito-QC (mCherry-GFP-FIS1) expressing cell lines. Cells were treated with 5 nM CAP-1902 for 16 h. Analysis was performed by counting the red puncta per cell, excluding the colocalizing signals between GFP and mCherry using the MetaXpress software. Leupeptin and Pepstatin A (LEUP) blockade allows for the calculation of the mitolysosomes flux as number of accumulated mitolysosomes overtime after leupeptin/pepstatin A addition. Representative images are shown in Figure S1H. Statistical analysis was performed by two-way ANOVA. (**D**) Representative high-throughput confocal micrographs of Ctrl and patient mutant fibroblasts labeled with GRP75 antibody (mitochondria - green), and DAPI (nuclei – blue). Cells were treated with 5 nM CAP-1902 for 16 h. Scale bars: 50 μm. **(E)** Quantification of mitochondrial mass as sum of area per cell from GRP75 images. Statistical analysis was performed by two-way ANOVA. **(F)** Relationship between the change in mitochondrial mass (GRP75 staining) and mitolysosomes flux (mito-QC expressing) in the fibroblasts treated with CAP-1902. **(G)** Quantification of mitolysosomes flux per cell as represented in H. Analysis was performed using the machine learning pixel classifier in AIVIA software. Leupeptin and Pepstatin A (LEUP) blockade allows for the calculation of the mitophagy flux as number of accumulated mitolysosomes overtime after leupeptin addition. Statistical analysis was performed by paired t-test. **(H)** Confocal micrographs of CIII-deficient (BCS1L) cells expressing mito-QC treated with 5 nM CAP-1902 for 16 h. Mito-QC was imaged using LSM880 Airyscan confocal microscopy. 10 μM Leupeptin/Pepstatin A (+LEUP) was used to block lysosomal degradation. Scale bars: 20 μm. **(I)** Segmentation of individualized mitochondria with low TMRE average intensity was used to count the individualized and depolarized mitochondria per cell shown in J. Statistical analysis was performed by one-way ANOVA. **(J)** Representative confocal micrographs of Ctrl, CIII-deficient cells treated with 5 nM CAP-1902 for 16h, stained with TMRE (red, top). MTG (see in Fig. S1K) was used as a mask to create the average TMRE intensity in MTG (heatmap blue:red, bottom). Scale bars: 20 μm. Cells were imaged with the LSM880 Airyscan microscope and analyzed by machine learning pixel classifier in AIVIA software. In all cases, data show mean ± SEM. Dots in graphs represent independent biological replicates.

Next, Ctrl and CV-deficient cells were used to assess mitochondrial matrix summed area as an indication of mitochondrial mass changes following treatment with a novel heteroaryl non-peptidic compound, CAP-1902 (Figure S1H). CAP-1902 mimics Ang-(1-7) and functions as a potent and selective MasR agonist. A 16 h treatment with CAP-1902 revealed that the highest concentration tested (500 nM) reduced mitochondrial mass in both Ctrl and ATP6 cells, whereas lower concentration (5, 50 and 100 nM) produced no detectable change.

To specifically study mitophagy flux, we stably expressed the mito-QC reporter in the mitochondrial disease fibroblasts (Figure 1C and S1I). Mito-QC encodes a tandem GFP-mCherry fluorescent protein attached to a mitochondria outer membrane target signal (amino acids 101-152) derived from the FIS1 anchor protein. Once a tagged mitochondrion is delivered to the lysosome, the GFP signal is quenched in the acidic environment, while the mCherry remains visible as a distinct red puncta.

The change in the number of red puncta corresponds to the number of mitochondria inside the autophagosomes that are fused to lysosomes, or mitolysosomes. In this case, since the basal number of mitolysosomes may not accurately reflect the mitophagy flux, that is, the total amount of mitolysosomes that a cell can make and degrade over time, a specific mitophagy flux calculation will be used when the compound is added. To accumulate mitolysosomes over time, inhibitors of lysosomal hydrolases and proteases, leupeptin and pepstatin A, were used to block degradation of mitolysosomes [23–25]. The ratio between the number of mitolysosomes per cell in presence and absence of the inhibitors was used for the calculation of mitolysosomes flux.

Because the treatment with Ang-(1-7) and CAP-1902 (500 nM) produced changes in mitochondrial mass suggestive of an increased mitochondrial turnover, we quantified both mitolysosomes flux in mito-QC expressing cells and mitochondrial mass in no mito-QC cells treated with CAP-1902 to more accurately assess mitochondrial turnover (Figure 1D-F). The addition of CAP-1902 at the lowest concentration tested (5 nM) improved mitolysosome flux in the CIII-deficient (P1:BCS1L^R73C/F368I^) and promoted a slight increase in Ctrl cells, while no difference was observed in CIII- (P2:BCS1L^R183C/R184C^) and CV- (ATP6^L156R^) deficient cells (Figure 1C-D). Interestingly, the increase in mitophagy flux in the CIII-deficient (P1:BCS1L) and Ctrl cells occurred without changes in mitochondrial mass (Figure 1E), which supports that CAP-1902 restored the balance in mitochondrial turnover (Figure 1F). No differences in mitochondrial morphological parameters were induced by CAP-1902 (Figure S1H-J).

Since previous data was produced using high-content screening microscopy, we repeated the mitolysosomes flux study using higher resolution confocal microscopy to validate our results and further investigate the mechanism of CAP-1902 on mitochondrial turnover. We focused on the CIII-deficient patient-derived fibroblast (P1:BCS1L), referred to as CIII henceforth. Detailed experiments performed on CIII-deficient fibroblasts confirmed the previously observed increase in mitolysosomes flux with CAP-1902 compared to untreated fibroblasts (Figure 1G-H).

Loss of mitochondrial membrane potential (ΔΨm) is a critical step in initiating mitophagy. Upon mitochondrial damage or dysfunction, the ΔΨm collapses, leading to the mitochondria either eventually recovering from the insult, or remaining individualized and depolarized to undergo mitophagy [26]. For that reason, it is important to understand whether CAP-1902 is causing depolarization or selectively acting on the turnover of depolarized mitochondria. Using the mitochondrial membrane potential-sensitive dye tetramethylrhodamine (TMRE) combined with MitoTracker green (MTG) staining (Figure 1I-J, S1L-N), we first measured the average intensity of TMRE in the MTG area. No differences were observed between Ctrl, CIII-deficient, and CAP-1902 treated cells. FCCP confirmed that TMRE/MTG staining was reporting on membrane potential with high sensitivity, both in Ctrl and CIII-deficient cells as expected (Figure S1L-N). This result indicates that the increase in mitophagy is not caused by CAP-1902 causing mitochondrial depolarization. To specifically measure selective mitophagy of the depolarized mitochondria, we analyzed the number of individualized and depolarized mitochondria in the cells. When individualized mitochondria, selected by MTG area, were counted, we observed a decrease in the number of small mitochondria after treatment with CAP-1902 (Figure S1N). Furthermore, by separating the individualized mitochondria and counting the number of small, depolarized organelles, we observed a decrease in the number of depolarized mitochondria in CAP-1902 treated cells (Figure 1I). These results indicate that CAP-1902 selectively targets the already depolarized mitochondria for degradation, instead of causing depolarization to induce mitophagy.

### Activation of the MasR by CAP-1902 increases mitochondrial biogenesis by restoring PGC1α translocation to the nucleus in CIII-deficient fibroblasts

To understand the mechanism by which CAP-1902 increases mitochondrial turnover, we performed RNAseq analysis in whole cells. We compared the differential transcriptional gene regulation between CIII-deficient untreated and treated with CAP-1902 for 24 h. Gene set enrichment analysis (GSEA) using the Reactome database with annotation of detailed mitochondrial processes (Figure S2A), together with a linked heat map of the leading-edge genes (Figure 2A), revealed a positive enrichment in respiratory electron transport and mitochondrial translation pathways induced by CAP-1902 treatment. In contrast, mitochondrial biogenesis pathways exhibited a bidirectional pattern, with both upregulated and downregulated genes (Figure 2A, S2A). Particularly, an increase in *PPRC1* (Figure 2A) is observed with CAP-1902 treatment of CIII-deficient cells. *PPRC1* encodes for PGC-1 Related Coactivator (PRC), which is shown to directly bind NRF1 (Nuclear Respiratory Factor 1) to promote mitochondrial biogenesis similarly to PGC1α (peroxisome proliferator-activated receptor gamma coactivator 1 alpha) [27]. Together with several of the upregulated transcripts in response to CAP-1902 related to electron transport chain (ETC) components, mitochondrial translation, and biogenesis (Figure 2A), these results suggest that the treatment promotes the renewal of mitochondrial proteins, by concurrently activating mitochondrial bioenergetic transcriptional programs and mitophagy.

**Figure 2.**
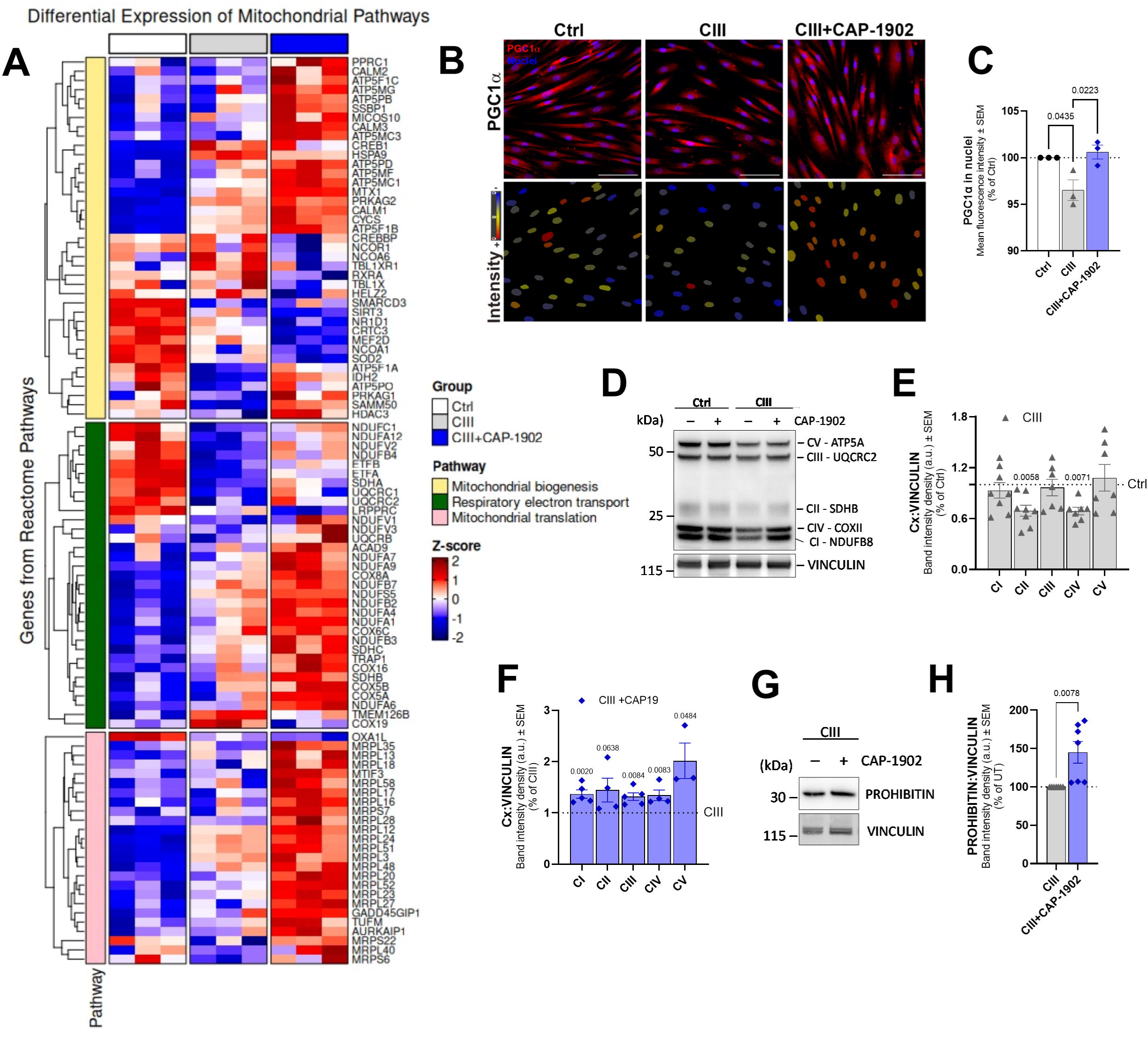
MasR agonist induces the nuclear translocation of PGC1α and increases expression of electron transport chain subunits in a CIII-deficiency model. **(A)** Heatmap of differentially expressed mitochondrial pathway genes across experimental conditions. The heatmap displays normalized expression (Z-scores) of genes within Reactome pathways: Respiratory Electron Transport, Mitochondrial Translation, and Mitochondrial Biogenesis; selected based on differential expression (p < 0.1) between drug-treated (CIII + CAP-1902) and untreated (CIII) cells. A control group (Ctrl) is included for baseline comparison. **(B**) Representative confocal micrographs of Ctrl, CIII-deficient untreated and treated with 5 nM CAP-1902 for 16 h labeled with PGC1α antibody. Scale bars: 100 μm. **(C)** Quantification of the average fluorescence intensity showing the distribution of PGC1α in the nuclei using the MetaExpress software. Statistical analysis was performed by one-way ANOVA. **(D)** Representative blots showing OXPHOS subunits from Ctrl, CIII-deficient and CIII-deficient treated with 5 nM CAP-1902 for 16 h. Vinculin was used as loading control. **(E)** Average protein expression of the OXPHOS subunits in Ctrl vs CIII-deficient cells. Statistical analysis was performed by unpaired t-test for each complex. (**F**) Average OXPHOS protein expression in CIII-deficient and CIII-deficient treated with 5 nM CAP-1902 for 16 h. Statistical analysis was performed by unpaired t-test for each complex. (**G**) Representative blots showing the expression of PROHIBITIN in CIII-deficient cells untreated and treated with 5 nM CAP-1902 for 16 h. Vinculin was used as loading control. (**H**) Average quantification of PROHIBITIN in CIII-deficient cells untreated and treated with 5 nM CAP-1902 for 16 h. Statistical analysis was performed by unpaired t-test. In all cases, data show mean ± SEM. Dots in graphs represent independent biological replicates.

To better explore the mitochondrial biogenesis pathway, we measured the translocation of PGC1α into the nuclei by immunofluorescence (Figure 2B-C). When comparing Ctrl cells to CIII-deficient cells, the latter showed a reduced amount of PGC1α in the nuclei that is restored after treatment with CAP-1902 (Figure 2C). Moreover, the amount of all the ETC components tested is increased with CAP-1902 treatment (Figure 2D-E), supporting our observations from the RNAseq analysis. CAP-1902 not only restored the levels of CII subunit (SDHB) and CIV subunit (COX2), decreased in CIII-deficient cells (Figure 2E), but also enhanced the protein levels of CI, CIII, and CV that were unchanged in the CIII-deficient cells (Figure 2F). The absence of changes in mitochondrial mass with CAP-1902 in CIII-deficient cells was confirmed by additional analysis of mitochondrial proteins from different subcompartments (Figure S2B-D). While in CIII-deficient compared to Ctrl cells, a decrease in MIC60 (inner membrane) and an increase in TOMM20 (outer membrane) were observed (Figure S2C), CAP-1902 treatment only increased levels of PROHIBITIN (Figure 2G) without affecting any of the additional mitochondrial markers tested (Figure S2D). Increased PROHIBITIN, together with the increase in ETC components with CAP-1902, suggested an increase in OXPHOS capacity per cristae, or alternatively, an increase in the number of cristae. Thus, to delve into the effects of CAP-1902 on cristae number, we used STED super-resolution microscopy to visualize cristae in live cells (Figure S2E-F). CAP-1902 showed no changes in the number of cristae, indicating that biogenesis induced by PGC1α is directed to renew OXPHOS components, increasing its capacity in the CIII-deficient cells.

### MAS receptor activation by CAP-1902 induces AMPK phosphorylation and increases ULK1/FUNDC1-dependent mitophagy

We next investigated the mechanism by which CAP-1902 is increasing PGC1α-mediated mitochondrial biogenesis. We first assessed AMP-activated protein kinase (AMPK)-mediated signaling, one of the signal transduction pathways for the translocation of PGC1α. The phosphorylation of AMPK at threonine 172 (T172) is a critical event in the activation of AMPK. This post-translational modification is a key regulatory mechanism that allows AMPK to sense cellular energy status and respond accordingly. CIII-deficient cells showed low levels of p-AMPK (T172) compared with Ctrl cells (Figure 3A-B), as expected when increased glycolysis to lactate is sufficient to preserve ADP/ATP ratio when mitochondria are deficient. However, CAP-1902 was still able to acutely activate p-AMPK (T172) after 10 and 30 minutes of treatment (Figure 3A, C), indicating a mechanism of AMPK-dependent PGC1α activation. As CAP-1902 is a small molecule that activates the MasR, we first verified the receptor’s presence in our cellular models. Indeed, MasR is expressed in both Ctrl and CIII-deficient cells but is not changed in the presence of CAP-1902 (Figure S3A-B). MasR antibody specificity was confirmed by using purified MasR tagged with GST as a positive control (Figure S3C). To validate whether the biological effects induced by CAP-1902 were indeed dependent on MasR activity, we tested whether the specific MasR inhibitor A779 [28] blocks CAP-1902 actions on AMPK phosphorylation. A779 treatment inhibited the activation of p-AMPK (T172), confirming that the rapid phosphorylation of AMPK is dependent on MasR activation by CAP-1902 (Figure 3D-E).

**Figure 3.**
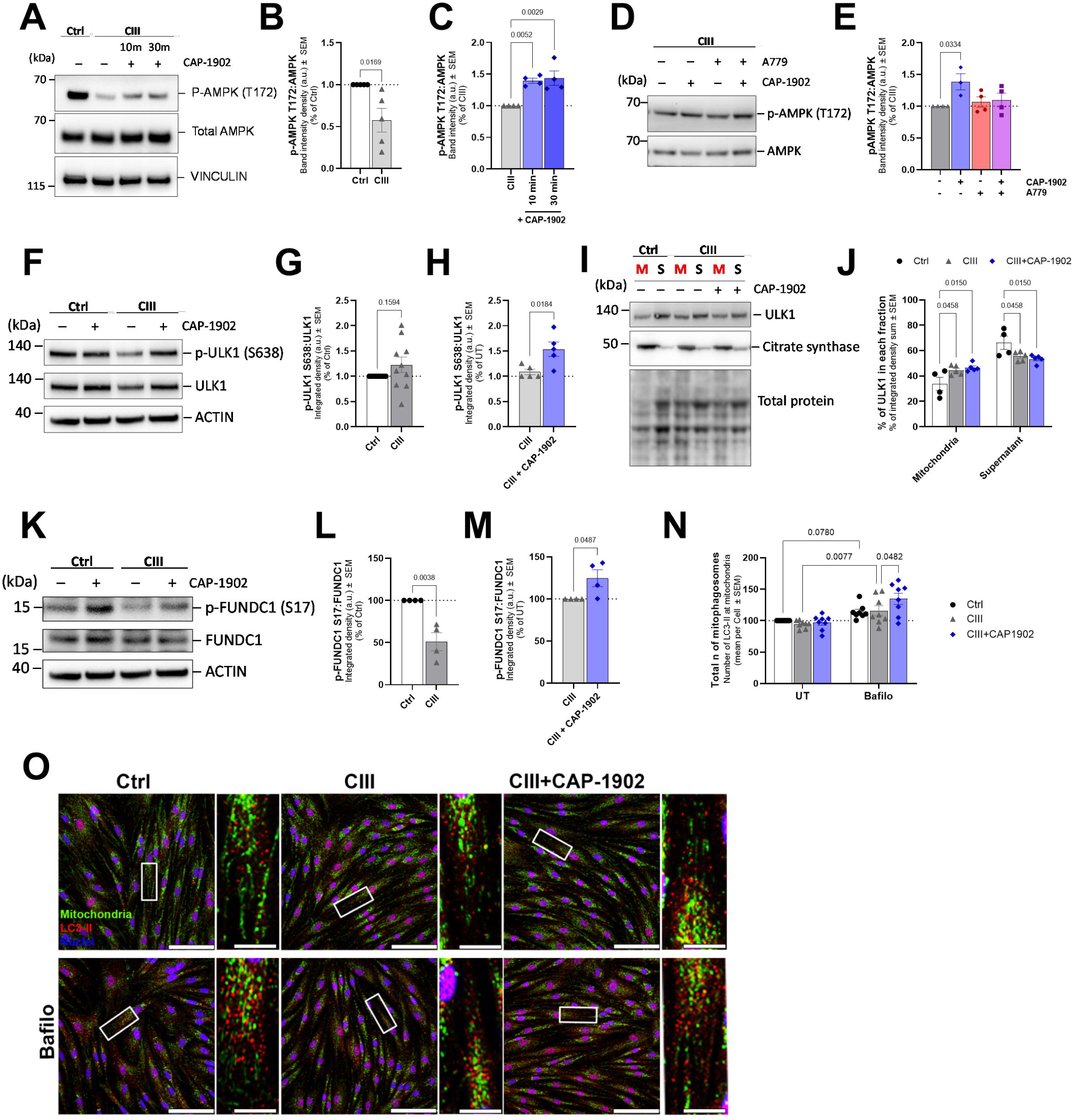
MasR activation by CAP-1902 triggers the AMPK/ULK1/FUNDC1 axis to increase mitophagy. **(A)** Representative blots of phosphorylated and total AMPK in Ctrl, CIII-deficient, and CIII-deficient cells treated with 5 nM CAP-1902 for 10 and 30 minutes. Vinculin was used as loading control. (**B**) Average expression of p-AMPK by total AMPK levels in Ctrl vs CIII-deficient, and (**C**) CIII-deficient untreated and treated with CAP-1902. Statistical analysis was performed using a t-test and one-way ANOVA, respectively. **(D)** Representative blots of phosphorylated and total AMPK in CIII-deficient untreated and treated with 1 μM A779 for 1h, plus and minus 5 nM CAP-1902 for the last 30 minutes. (**E**) Average expression of p-AMPK by total AMPK levels. Statistical analysis was performed by one-way ANOVA. **(F)** Representative blots of phosphorylated (S638) ULK1 and total ULK1 in Ctrl, CIII-deficient, and CIII-deficient cells treated with 5 nM CAP-1902 for 16 h. ACTIN was used as loading control. **(G)** Average expression of p-ULK1 by total ULK1 levels in Ctrl vs CIII-deficient untreated and (**H)** CIII-deficient untreated and treated with CAP-1902. Statistical analysis was performed by unpaired t-test. **(I)** Representative blots of ULK1 in mitochondria-enriched fractions (M) vs the corresponding supernatant (S) in Ctrl, CIII-deficient, and CIII-deficient cells treated with 5 nM CAP-1902 for 16 h. Citrate synthase is included as a control for mitochondrial enrichment, and total protein stain as loading control. **(J)** Quantification of the expression of ULK1/total protein in each fraction. Statistical analysis was performed by two-way ANOVA. **(K)** Representative blots of phosphorylated (S17) FUNDC1 and total FUNDC1 in Ctrl, CIII-deficient, and CIII-deficient cells treated with 5 nM CAP-1902 for 16 h. ACTIN was used as loading control. **(L)** Average expression of p-FUNDC1 by total FUNDC1 levels in Ctrl vs CIII-deficient untreated and **(M)** CIII-deficient untreated and treated with CAP-1902. Statistical analysis was performed by unpaired t-test. (**N**) Number of mitolysosomes in Ctrl, CIII-deficient, and CIII-deficient cells treated with 5 nM CAP-1902 for 16 h. Quantification represents the average number of LC3-II puncta touching mitochondria (mitophagosomes) per cell. Statistical analysis was performed by two-way ANOVA. (**O**) Confocal micrographs of Ctrl, CIII, and CIII cells treated with 5 nM CAP-1902 ± bafilomycin for 16 h, labeled with LC3B (autophagosome – red), GRP75 (mitochondria - green), and DAPI (nuclei - blue). Scale bars: 100 μm and 5 μm in the zoomed image. In all cases, data show mean ± SEM. Dots in graphs represent independent biological replicates.

AMPK also has a role increasing mitophagy that it is independent of PGC1α. AMPK directly phosphorylates ULK1 on at least four residues in mouse models: Ser467, Ser555, Thr574, and Ser637, which leads to an increase in mitophagosome formation [29]. We checked the phosphorylation of human ULK1 at Ser638, corresponding to S637 in rodents, in CIII-deficient cells, finding that pULK1 Ser638 levels were not changed (Figure 3F-G). However, the amount of ULK1 protein was increased in mitochondrial fractions from CIII-deficient fibroblasts (Figure 3I-J), suggesting an adaptation to defective ULK1 activity promoting mitophagy. Remarkably, CAP-1902 treatment induced an increase in the phosphorylation of ULK1 at Ser638, without changing the total amount of ULK1 in the CIII-deficient cells (Figure 3F, H).

Activated ULK1 induces mitophagy by phosphorylating the FUN14 domain-containing protein 1 (FUNDC1) mitochondrial cargo receptor [30]. FUNDC1 is phosphorylated by ULK1 on Ser17 close to its characteristic microtubule-associated protein 1A/1B light chain 3B (LC3)-interacting region domain (LIR domain), which helps to promote its interaction with LC3 and start mitophagy [30]. To confirm whether FUNDC1 phosphorylation was increased by CAP-1902 treatment, as expected from ULK1 activation, we measured the phosphorylation of FUNDC1 at Ser17 in the CIII-deficient cells (Figure 3K-L). While p-FUNDC1 is decreased in CIII-deficient compared to Ctrl cells, the treatment with CAP-1902 can revert this phenotype increasing the phosphorylation of FUNDC1 (Figure 3K, M), and therefore explaining the previously observed increased mitophagy flux. Additionally, CAP-1902 increases FUNDC1 levels in mitochondrial-enriched fractions, while no differences were observed between CIII-deficient and Ctrl cells (Figure S3D-F).

Specifically, Ser17 phosphorylation as well as Ser13 and Tyr18 dephosphorylation of the FUNDC1 sequence can enhance its interaction with LC3 and the recruitment of autophagosomes to the dysfunctional mitochondria to form the mitophagosome [31]. To specifically measure the formation of the mitophagosome, Ctrl and CIII-deficient cells were co-stained with anti-LC3-II antibody and a mitochondrial marker. Autophagic vesicles were imaged, and mitophagosomes defined as the vesicles touching the mitochondria (Figure 3O). Bafilomycin was used to block autophagic degradation to assess the flux of mitophagosomes. CIII-deficient cells showed no differences in the flux of mitophagosomes compared with Ctrl cells (Figure 3N-O), while the treatment with CAP-1902 increased the flux of mitophagosomes in CIII-deficient cells (Figure 3N). Western blotting analyses of LC3-II in mitochondrial enriched fractions further confirmed the increase in the engulfment of mitochondria into autophagosome induced by CAP-1902 treatment (Figure S3G-H). These results indicate that CAP-1902 induced phosphorylation of Ser17 in FUNDC1 can facilitate its interaction with LC3-II to form the mitophagosome, providing a mechanism for the increase in mitophagic flux observed (Figure 1G). Together our data suggest that the activation of MasR by CAP-1902 leads to the downstream activation of the AMPK/ULK1/FUNDC1 signaling pathway activating mitophagy and increasing the flux of mitophagosomes and mitolysosomes in CIII-deficient cells.

Validation of AMPK-dependent activation of mitophagy by CAP-1902 was pursued by blocking AMPK activation with Dorsomorphin (Figure S4A-B). Dorsomorphin inhibits AMPK by competitively blocking its ATP-binding site, preventing activation by upstream kinases and downstream signaling. Decreased activation of AMPK and ACC by phosphorylation was observed after a 30 minutes treatment with 10 μM Dorsomorphin in Ctrl cells, and a slight decrease was observed in CIII-deficient cells with 2.5 μM of Dorsomorphin. The difference in concentration needed to decrease AMPK activation in these cells may be due to the low levels of AMPK present in CIII-deficient cells. To verify if Dorsomorphin could be used to block CAP-1902 induction of mitophagy, we tested the ability of the cells to survive a long treatment with the AMPK inhibitor and observed decreased cell viability after 24 h of treatment (Figure S4C-D). This result abolished our ability to directly test if mitophagy induced by CAP-1902 is AMPK-dependent.

### FUNDC1 is required for the formation of the mitophagosome induced by MasR activation

Our results indicated that in CAP-1902 treated cells, FUNDC1 activation facilitates its interaction with LC3-II to form the mitophagosome. To confirm that FUNDC1 is the main player increasing mitophagy in our model, we knocked down FUNDC1 in CIII-deficient cells by shRNA lentiviral transduction (CIII^FUNDC1_KD^). GFP expression integrated in the lentiviral transduction confirmed transduction (Figure 4H), while knockdown efficiency was confirmed by western blot of FUNDC1 (Figure S4E). Decreased expression of FUNDC1 in CIII-deficient cells caused a major decrease in autophagosome degradation, as observed by the increased amount of mitophagosomes that was insensitive to bafilomycin treatment (Figure 4A-B). This result indicates a major dependency of autophagosome and mitophagosome clearance on FUNDC1 in CIII-deficient cells, which was unexpected as FUNCD1 affects autophagosome formation, not clearance. In this case, knocking down FUNDC1 blocked the effect of CAP-1902 in increasing flux of mitophagosomes (Figure 4A, C), but given the effects on clearance, we cannot unequivocally conclude whether this blockage reflects just an inability of CAP-1902 to increase mitophagosome formation.

**Figure 4.**
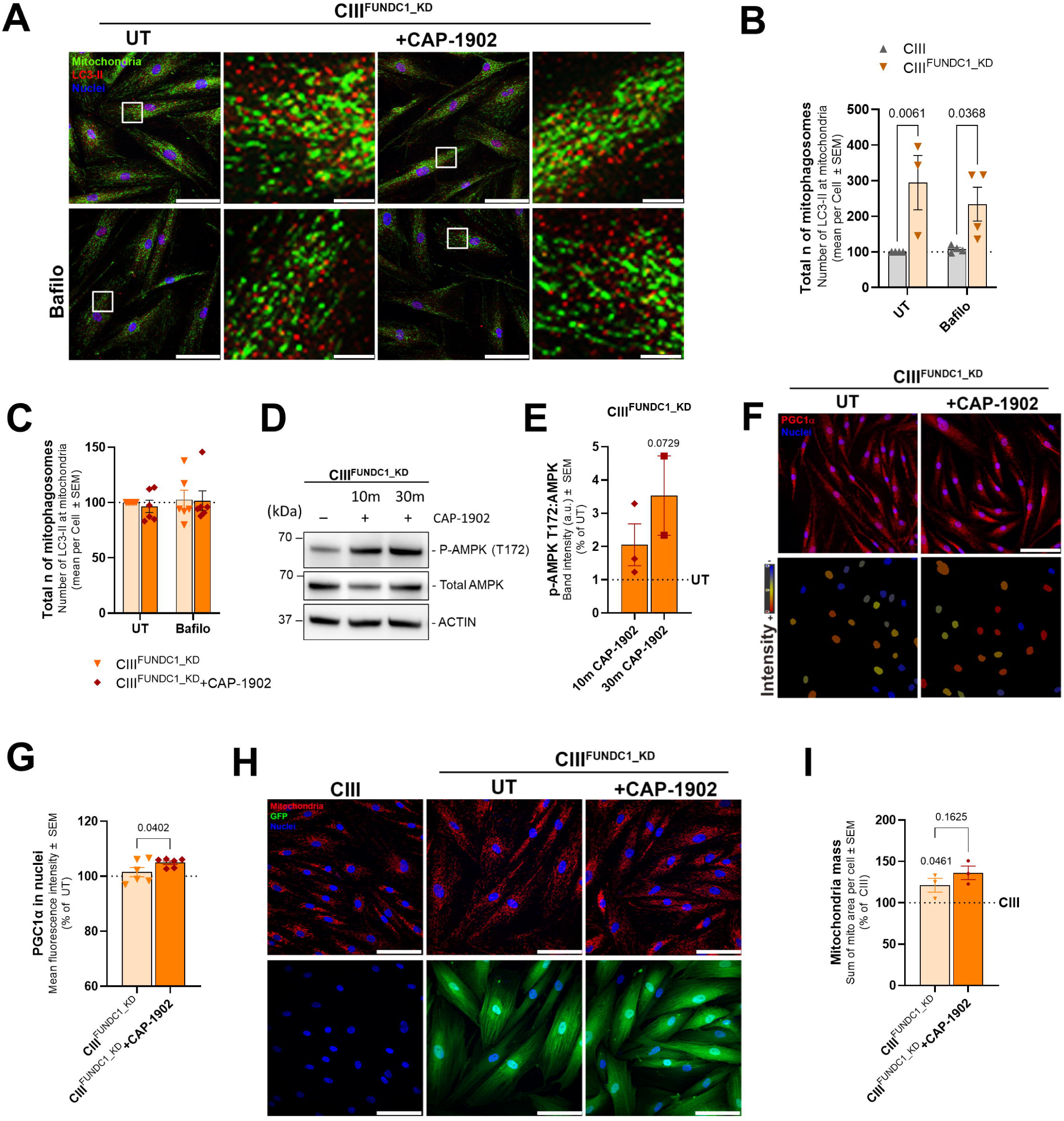
FUNDC1 knockdown abolishes MasR induced mitophagy. **(A)** Representative high-throughput confocal micrographs of CIII-deficient cells with shRNA knockdown for FUNDC1 (CIII^FUNDC1_KD^) untreated and treated with 5 nM CAP-1902 for 16h. Autophagy flux is blocked using 100 nM bafilomycin for 16 h. Scale bars: 100 μm and 5 μm in the zoomed image. (**B**) Number of mitolysosomes in CIII-deficient and CIII^FUNDC1_KD^. Quantification represents the average number of LC3-II puncta touching mitochondria (mitophagosomes) per cell. Statistical analysis was performed by two-way ANOVA. (**C**) Number of mitophagosomes in CIII^FUNDC1_KD^ untreated and treated with 5 nM CAP-1902 for 16h. Quantification represents the average of LC3-II puncta at mitochondria per cell. Statistical analysis was performed by two-way ANOVA. (**D**) Representative blots of phosphorylated and total AMPK in CIII^FUNDC1_KD^ untreated and treated with 5 nM CAP-1902 for 10 and 30 minutes. ACTIN was used as loading control. (**E**) Average expression of p-AMPK referred to total AMPK in the conditions specified in D. (**F**) Representative confocal micrographs of CIII^FUNDC1_KD^ untreated and treated with 5nM CAP-1902 for 16 h labeled with PGC1α antibody (red) and DAPI (nuclei – blue) (top). Mask showing the intensity of the nuclei in the previous images (bottom). Scale bars: 100 μm. (**G**) Quantification of the average PGC1α fluorescence intensity in the nuclei. Statistical analysis was performed by paired t-test. (**H**) Representative confocal micrographs of CIII-deficient, CIII^FUNDC1_KD^ cells untreated and treated with 5 nM CAP-1902 for 16 h labeled with GRP75 antibody (mitochondria – red) and DAPI (nuclei – blue) (top). GFP signal from the same samples indicates infection with FUNDC1 shRNA (bottom). Scale bars: 100 μm. (**I**) Quantification of mitochondrial mass in CIII^FUNDC1_KD^ untreated and treated with 5 nM CAP-1902 for 16 h compared to CIII-deficient cells (100%). Statistical analysis was performed by paired t-test. In all cases, data show mean ± SEM. Dots in graphs represent independent biological replicates.

Next, we tested if the effect of CAP-1902 on AMPK and PGC1α were impacted by FUNDC1 downregulation. Phosphorylation of AMPK by CAP-1902 was still observed (Figure 4D-E), as well as the increase on PGC1α nuclear translocation in CIII^FUNDC1_KD^ treated cells (Figure 4F-G). An increase in mitochondrial mass was observed in the CIII^FUNDC1_KD^ compared with CIII-deficient cells (Fig. 4H-I), which we attributed to the decrease in FUNDC1-dependent mitophagosome clearance. In this context, while CAP-1902 still increased PGC1α translocation to the nucleus in CIII^FUNDC1_KD^ cells, no further increase in mitochondrial mass is observed (Figure 4H-I). This plateau effect is likely due to stress-induced mechanisms that suppress translation of PGC1α-responsive transcripts when mitophagy is compromised.

### CAP-1902 restores lysosomal distribution in CIII-deficient fibroblasts

Mitochondrial depolarization can trigger both receptor-mediated and ubiquitin-dependent mitophagy by targeting the dysfunctional mitochondria for degradation. All these targeted mitochondria will be delivered to autophagosomes (mitophagosomes) that will commonly be fused to the lysosome, forming the mitolysosome in preparation for degradation. To check if the activation of MasR by CAP-1902 also targets lysosomal pathways, we first measured levels of the Lysosome-Associated Membrane Protein 1 (LAMP1) by western blotting and confocal microscopy (Figure S5A-D). No differences were observed in the levels of LAMP1 protein or intensity in the cytoplasm of CIII-deficient compared to Ctrl cells. Treatment with CAP-1902 did not change levels of LAMP1, indicating no increase in the number of lysosomes (Figure S5C-D). While no differences were observed in the amount of LAMP1, a change in the distribution pattern of the LAMP1 vesicles (lysosomes) was observed (Figure 5A-B). Lysosome retrograde transport and perinuclear clustering have been shown to activate autophagy, while the anterograde lysosome movement toward the cell periphery is linked to the inhibition of autophagy [32, 33]. Knowing that, we determined the intracellular distribution of LAMP1 vesicles, in which we categorized the vesicles according to their distance from the nucleus and assessed the number of peripheral lysosomes. In CIII-deficient cells, more peripheral LAMP1 vesicles are observed than in Ctrl cells (Figure 5B), corresponding to an inhibition of autophagy. This was rescued by CAP-1902, indicating that lysosomal distribution has a role in the CAP-1902 mechanism of action. Next, we evaluated whether lysosomal function is changed after CAP-1902 treatment by using DQ-BSA colocalization with LAMP1 vesicles (Figure S5E-F). This technique relies on the cleavage of the self-quenched DQ™ Red BSA protease substrates in an acidic compartment, such as the lysosome, to generate a highly fluorescent product [34]. If an increase in fluorescence intensity is observed, more active proteases are present in the lysosomal compartment. CAP-1902 treatment did not increase fluorescence activity in CIII-deficient cells (Figure S5F). Indeed, lysosomal activity was increased in CIII-deficient cells when compared with Ctrl cells, which demonstrated that the defective distribution of lysosomes, rather than lysosomal activity itself, was the only lysosomal defect induced by lack of CIII activity. Altogether, these results indicate that CAP-1902 restores the distribution of lysosomes without increasing lysosomal biogenesis and activity.

**Figure 5.**
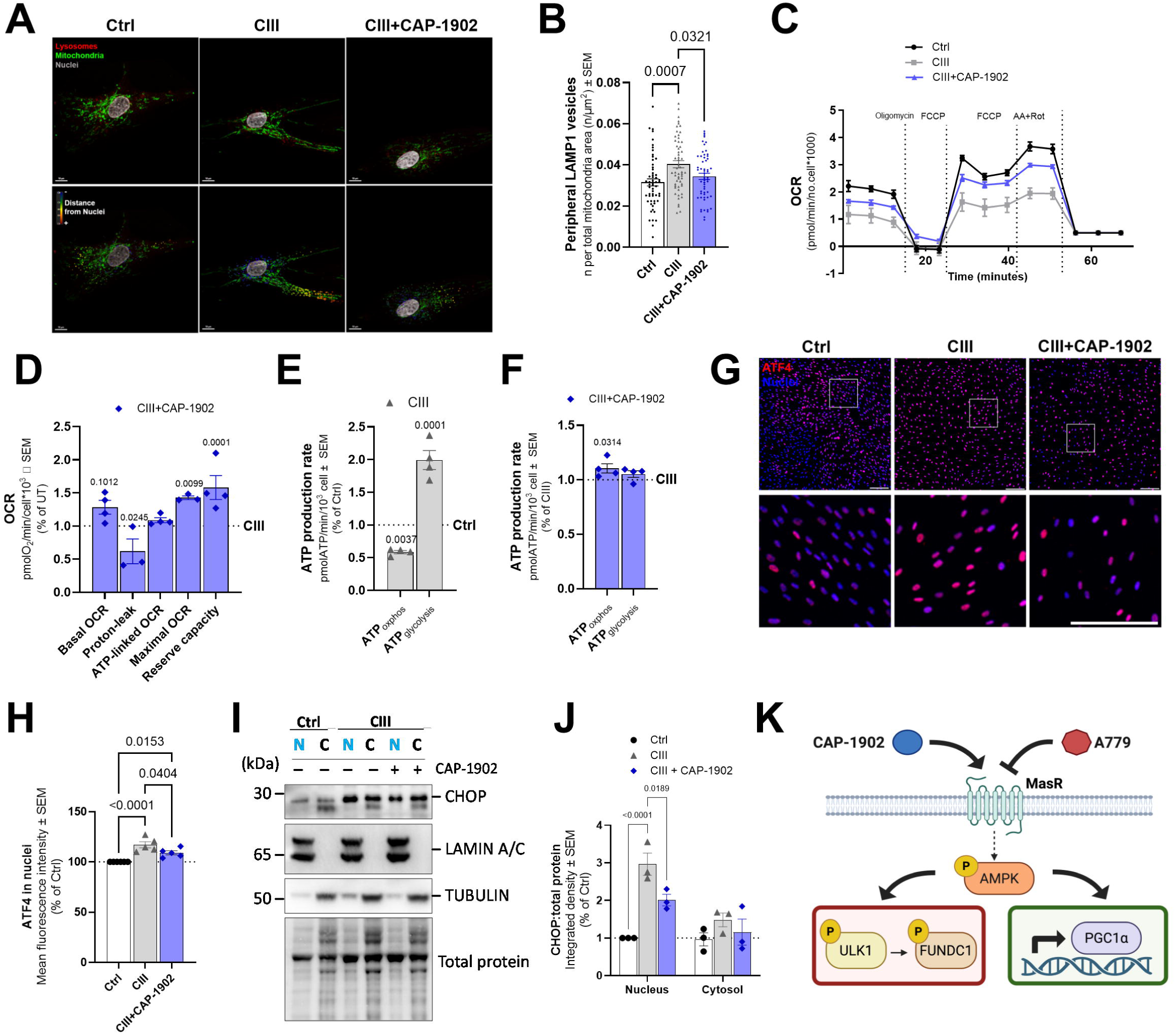
MasR agonist improves mitochondrial OXPHOS and lysosomal distribution, while decreasing the activity of the Integrated Stress Response effectors ATF4 and CHOP. **(A)** Confocal micrographs of Ctrl, CIII-deficient, and CIII-deficient cells treated with 5 nM CAP-1902 for 16 h, labeled with LAMP1 (lysosomes - red), TOMM20 (mitochondria - green), and DAPI (grey). Scale bars: 10 μm. The distance of each lysosome from the nuclei is highlighted (heatmap blue:red). **(B)** Quantification of LAMP1 vesicles in the cellular periphery (above 75 percentile distance from the nuclei) represented as the number of vesicles per mitochondrial area. Statistical analysis was performed by one-way ANOVA. **(C)** Representative oxygen consumption traces from a Mito Stress test assay in Ctrl, CIII-deficient, and CIII-deficient cells treated with 5 nM CAP-1902 for 24 h. **(D)** Respiratory parameters in CIII-deficient compared to CIII-deficient cells treated with 5 nM CAP-1902 for 24 h. Statistical analysis was performed by two-way ANOVA. **(E)** ATP production rate calculations in Ctrl compared to CIII-deficient cells. Statistical analysis was performed by two-way ANOVA. **(F)** ATP production rate calculations in CIII-deficient compared to CIII-deficient cells treated with 5 nM CAP-1902 for 24 h. Statistical analysis was performed by two-way ANOVA. (**G**) Representative confocal micrographs of Ctrl, CIII-deficient, and CIII-deficient cells treated with 5 nM CAP-1902 for 16 h, labeled with ATF4 antibody (red) and DAPI (blue). Cells were imaged using high-throughput confocal microscopy and analyzed using MetaXpress software. Scale bars: 100 μm. DAPI was used to create a nuclear mask and the average fluorescence intensity was measured in the nuclei. (**H**) Quantification of ATF4 intensity in the nuclei. Statistical analysis was performed by one-way ANOVA. **(I)** Representative blots of CHOP in nuclear (N) versus cytosolic (C) fractions in Ctrl, CIII-deficient, and CIII-deficient cells treated with 5 nM CAP-1902 for 24 h. LAMIN A/C is included as a control for nuclei, TUBULIN as a control for cytosol, and toal protein stain as loading control. (**J**) Quantification of the expression of CHOP by total protein in both fractions. Statistical analysis was performed by two-way ANOVA. (**K**) Schematic representing the proposed signaling pathway of CAP-1902. In all cases, data show mean ± SEM. Dots in graphs represent independent biological replicates.

### CAP-1902 improves respiration and decreases markers of integrated stress response (ISR) in CIII-deficient fibroblasts

The results above show increased mitochondrial turnover, suggesting that CIII-deficient cells under CAP-1902 treatment should experience an improvement in mitochondrial function. To investigate this, we examined the respiratory function of the CIII-deficient cells compared with Ctrl cells by measuring oxygen consumption (OCR) and extracellular acidification (ECAR) (Figure 5C-F; Figure S5G-K). When compared with Ctrl cells, CIII-deficient cells showed a decrease in basal, ATP-linked, and maximal respiration, as well as a compromised reserve capacity in glucose media (Figure 5C, S5G). Proton leak is however increased, indicating inefficient respiration. This deficient respiratory phenotype correlates with increased ECAR levels in the CIII-deficient mutant (Figure S5H), as expected from an increase in glycolysis to compensate for a deficiency in OXPHOS. Treatment with CAP-1902 for 24 h rescues maximal respiration and reserve capacity, while decreasing proton-leak (Figure 5C-D), all of which is indicative of a healthier mitochondrial population. Additionally, when calculating the ATP production rate derived from OXPHOS we observed a rescue of this parameter in CIII-deficient cells treated with CAP-1902 (Figure 5E-F). When respirometry was performed in galactose, where cells are forced to rely on OXPHOS to produce ATP, the impaired respiratory phenotype remains in CIII-deficient cells, showing an increase in proton leak and a decrease in maximal respiration (Figure S5J). Rescue of both proton-leak and maximal OCR was observed with CAP-1902 treatment under galactose as well (Figure S5K).

Additionally, we tested a few players in the cellular stress response [35]. In mitochondrial diseases, Activating Transcription Factor 4 (ATF4) is activated and translocated to nuclei to help coordinate the expression of several genes associated with the integrated stress response (ISR), which is a complex network of signaling pathways that helps cells adapt to stress conditions. To confirm the translocation of ATF4 to the nuclei, we measured the intensity of the ATF4 antibody staining in the nuclei of Ctrl and CIII-deficient cells (Figure 5G-H). We observed an increase in levels of ATF4 in the nuclei of CIII-deficient cells, which decreased with CAP-1902, suggesting that restoration of OXPHOS diminished the activation of ISR.

One downstream target of ATF4 is the C/EBP Homologous Protein (CHOP) [25]. ATF4 binds to the C/EBP-ATF response element in the CHOP promoter region, leading to increased CHOP transcription. CHOP is a proapoptotic transcription factor, and its induction is associated with prolonged or severe ER stress [36]. While ATF4 helps cells adapt to stress, CHOP has a role in triggering apoptosis under conditions where cellular recovery is not possible. Next, we measured the levels of CHOP in cytosolic and nuclear fractions to determine the degree of ATF4 activation induced by CIII-deficiency (Figure 5I-J). CHOP is increased in the nuclear fraction of CIII-deficient cells and decreased with CAP-1902 treatment, indicating the mitigation of prolonged ISR activation induced by CIII deficiency.

Overall, our results show that in a model of CIII deficiency, CAP 1902 reduces cellular stress and improves bioenergetics by activating MasR, which triggers AMPK phosphorylation and drives mitochondrial turnover through the ULK1–FUNDC1 axis, while also activating PGC1α to enhance mitochondrial biogenesis (Figure 5K).

## DISCUSSION

Mitochondrial function is essential to metabolic homeostasis, and mitochondrial quality control is a crucial aspect to maintain a healthy and functional mitochondria population. In this way, cellular homeostasis depends on the achievement of an appropriate balance between mitochondrial biogenesis and mitophagy, or mitochondrial turnover [37, 38]. In this study, we have demonstrated that CAP-1902, a novel agonist of MasR, can induce mitochondrial turnover in a CIII-deficiency model of primary mitochondrial disease, in which dysfunctional mitochondria are accumulated, and turnover is impaired. We further show the benefits to (1) mitochondrial function by the reduction in depolarized mitochondria and improved bioenergetics, and (2) overall cellular health by correction of lysosomal distribution and decreased activation of the ISR components ATF4 and CHOP.

In the CIII-deficient cells, we have shown that CAP-1902 can increase mitophagy flux by restoring the number of mitolysosomes and mitophagosomes over time. Remarkably, damaged and depolarized mitochondria are selectively targeted in this enhanced mitophagy, indicating a potential therapy for mitochondrial diseases. Recent studies support the therapeutic potential of modulating mitophagy in mitochondrial diseases [2–6]. Some natural compounds (such as urolithin A) and other interventions like AICAR (an AMP analog) or coenzyme Q (CoQ) can activate mitophagy and restore homeostasis in MELAS and aging models [39, 40]. Rapamycin is the most extensively studied drug in this context and has been shown to restore mitophagy across various cellular and animal models, acting through mTOR-dependent regulation of mitochondrial recycling [2, 3, 41]. While rapamycin shows therapeutic promise, its potential adverse effects, particularly on immune function and protein synthesis, pose limitations for long-term use [42]. Therefore, identifying alternative mitophagy inducers with fewer chronic side effects is a high priority in the effort to halt or slow the progression of mitochondrial diseases. Notably, activation of the MasR by Ang-(1-7) has been shown to reduce neuroinflammation and cognitive decline in Alzheimer’s disease models, in part by decreasing amyloid-β and phospho-tau accumulation, a process that could be mediated by enhanced autophagy [43]. In other models of cerebral ischemia and aneurysm, MasR activation promoted angiogenesis and neuroprotection by reducing neuronal cell death [44].

In parallel to the mitophagy activation, CAP-1902 can induce mitochondrial biogenesis as evidenced by the upregulation of multiple transcripts and increased expression of proteins related to mitochondria renewal and the increased translocation of PGC1α to the nucleus. The lack of change in mitochondrial mass observed with CAP-1902 treatment despite the activation of such a transcriptional program, highlights that the activation of mitochondria degradation and biogenesis by CAP-1902 are balanced, illustrating the uniqueness of dual CAP-1902 action in mitochondria. Drugs such as mito-CP and mito-metformin have been shown to induce mitophagy, but also to inhibit ATP-linked oxygen consumption and reduce maximal respiratory capacity [46], potentially due to excessive removal of functional mitochondria. Another example is that promoting mitochondrial biogenesis alone with LY344864, an HTR1F agonist, transiently enhances mitochondrial oxygen consumption. However, this effect is transient as mitophagy then restores mitochondrial mass density [47]. Interestingly, treatment with Urolithin A, a mitophagy inducer, led to reduced basal respiration in *C. elegans* and C2C12 myotubes, despite preserved maximal respiratory capacity. The authors attributed this effect to an indirect inhibition of Complex I [40], showing the importance of understanding the drug’s mechanism of action. An inhibition in CI can be particularly detrimental in the context of mitochondrial diseases characterized by ETC deficiencies. In our results, the balanced mitochondrial turnover achieved by CAP-1902 was able to improve mitochondrial respiration. Confirming such balanced effects, CAP-1902 treatment rescued ATP production derived from oxidative phosphorylation in CIII-deficient cells, without changing mitochondrial mass.

Within our study, we explored the mechanism by which CAP-1902 induces mitochondrial turnover. Following CAP-1902 binding to MasR, we observe rapid AMPK activation that was blocked by A779, a MasR antagonist [28]. This supports MasR involvement but does not exclude indirect activation of AMPK as the proximate trigger. Attempts to inhibit AMPK with Dorsomorphin caused reduced cell survival and therefore hindered our ability to confirm pathway specificity. AMPK is a major regulator of cellular and mitochondrial bioenergetic homeostasis. AMPK coordinates and regulates eukaryotic mitophagy and mitochondrial biogenesis, making it a critical player and target in drug development [21]. For example, treatments with AICAR and Coenzyme Q10 have been shown to induce mitophagy and increase mitochondrial mass while also activating AMPK [39, 48]. Several stimuli can activate AMPK, including the activation of the protective arm of the RAS [22], and in this work, the activation of the MasR. Downstream of AMPK, CAP-1902 activates ULK1, an inducer of autophagy in mammals. This connection between AMPK and ULK1 is well established in the literature [19, 20, 49] and has been previously reported for other mitophagy-inducing compounds [39]. Recent studies indicate that AMPK can exert opposing effects on mitophagy depending on the cellular context, by either inhibiting NIX-dependent mitophagy through ULK1 phosphorylation or activating ULK1-driven mitophagy [50]. In line with our findings that CAP-1902 stimulates AMPK-ULK1 signaling to promote mitophagy in fibroblasts, the authors show that MK-8722 similarly activates the AMPK–ULK1 phosphorylation cascade in skeletal muscle and induces mitophagy, reinforcing that AMPK’s impact on mitophagy is both tissue- and pathway-specific.

Followed by the activation of ULK1 by CAP-1902, the mitophagy receptor FUNDC1 is phosphorylated at Ser17, promoting its translocation to the mitochondria. FUNDC1 has been described to regulate mitophagy by the direct interaction of Ser17 and LC3-II and it is currently being studied as a therapeutic target for the prevention or treatment of diseases like cardiovascular disease, metabolic syndrome, pulmonary hypertension, and neurodegenerative diseases [51–53]. Phosphorylation of Ser17 recruits LC3-II to the mitochondria to form the mitophagosome. CAP-1902 increased the mitophagosome flux and knocking down FUNDC1 was shown to inhibit this effect. It has also been reported that FUNDC1-dependent mitophagy is directly coupled with mitochondrial biogenesis through the PGC1α/NRF1 pathway [54]. In our work, the activation of PGC1α by CAP-1902 addition was not blocked by knocking down FUNDC1 but limited mitochondrial mass increase was observed. These results indicate that the activation of PGC1α by CAP-1902 happens upstream of FUNDC1, but effective mitochondrial biogenesis can be further affected by the downregulation of FUNDC1. As CAP-1902 activation of AMPK happens as fast as in 30 minutes after addition to the media, regulation of PGC1α, a well-described regulator of mitochondrial biogenesis by direct activation of AMPK, could occur earlier than ULK1/FUNDC1 activation [22, 55, 56]. These interactions highlight that the activation of the AMPK/ULK1/FUNDC1 signaling pathway by CAP-1902 is mediating the coordination between synthesis and degradation of mitochondria improving mitochondrial homeostasis in CIII-deficient cells.

Our study has described the first potential GPCR-mediated treatment of mitochondrial diseases by inducing mitochondrial turnover. Importantly, this is the first study where MasR, a GPCR and component of the protective arm of RAS, can impact mitochondrial function and potentially serve as a therapeutic target for mitochondrial disorders. Recently, semaglutide use in tauopathy models showed improvement in cognitive regression and increased levels of Ang-(1-7). The activation of the protective arm of RAS was shown to promote AMPK/GSK-3B and Sirt-1/FOXO1 induced autophagic maturation [57]. Moreover, BIO101 (Sarconeos), a small-molecule activator of MasR with anabolic and pro-differentiating effects on muscle cells, has advanced into Phase 3 clinical program (SARA-31) for sarcopenia, an age-related progressive loss of skeletal muscle mass and function [45]. The clinical development of a MasR agonist for muscle degeneration underscores the therapeutic potential of targeting the protective RAS arm and further supports the concept that MasR may enhance mitochondrial quality control in diseases characterized by mitochondrial dysfunction.

Together with our study, this demonstrates that MasR, and the protective arm of RAS, can be exploited to modulate autophagy. In our study, we found that in the CIII-deficient cellular model, CAP-1902 mechanism of action is exerted by the AMPK/ULK1/FUNDC1 and AMPK/PGC1α signaling pathway. Moreover, this is the first study to link MasR to the mitophagy receptor, FUNDC1, and further associate a mitophagy activator with a particular mitophagy receptor. These findings have opened the RAS pathway to exploration in mitochondrial quality control and underscore the importance of achieving a balanced mitochondrial turnover by potential therapeutics developed for mitochondrial diseases.

## METHODS

### Cell culture

Primary fibroblasts derived from patients were acquired from Telethon Network Genetic Biobank (P1:BCS1L^R73C/F368I^, P2:BCS1L^R183C/R184C^), Coriell Institute for Medical Research (ATP6^L156R^), and Control. Cells were maintained in Eagle’s Minimum Essential Medium (EMEM; ATCC, 30-2003) supplemented with 15% FBS (Gibco, 6000-069) and antibiotic–antimycotic (Gibco, 15240-062) in a humidified atmosphere at 37°C and 5% CO2. Lentiviral infection of pLenti-mito-QC (produced by Welgen, Inc) and pGFP-C-shLenti-FUNDC1 (Origene, TL304445V) was performed using 8 mg/mL polybrene (Sigma-Aldrich, TR1003) in EMEM for 30 h and positive selection was performed by 1 μg/mL puromycin (Gibco, A1113803).

### Treatments

In all cases, except for experiments involving Angiotensin 1-7, cells were preconditioned for 24 h in phenol red-free Dulbecco’s Modified Eagle Medium (DMEM; Thermo Fisher Scientific, A1443001) containing 4.5 g/L glucose (Sigma-Aldrich, G5767) supplemented with 2 mM glutamine (Gibco, 25030-081), 1 mM sodium pyruvate (Gibco, 11360-070), 1% FBS (Gibco, 16000-069), and antibiotic–antimycotic (Gibco, 15240062) before adding the treatment. In experiments involving Angiotensin 1-7 (MedChem Express, HY-12403), the same phenol red-free media was used but without any FBS added. Additional compounds used in the study were: A779 (MedChem Express, HY-P0216), Bafilomycin A1 (SelleckChem, S1413), Leupeptin (Sigma-Aldrich, L2884), Pepstatin A (Sigma-Aldrich, P5318), Dorsomorphin Dihydrocholride (Tocris, 3093). These compounds were prepared in the solvent specified by the manufacturer and added directly to the media at the concentration and time specified in the figures. CAP-1902 compound was obtained from Capacity Bio and used at 5 nM in all the experiments. Treatment time is 16 h unless otherwise specified.

### Immunofluorescence

Cells were fixed with 4% paraformaldehyde (Thermo Fisher Scientific, 043368-9M) in phosphate-buffered saline (PBS; Gibco, 14190-144) for 20 min at RT, permeabilized (0.1% Triton X-100 (Sigma-Aldrich, T8787), 0.05% sodium deoxycholate (Sigma-Aldrich, 30970) in PBS), blocked, and stained with primary and secondary antibodies in blocking solution (5% donkey serum (Millipore Sigma, S30-M)). Coverslips were mounted with Mowiol. Image acquisition was performed using either a Zeiss LSM880 Confocal system with Airyscan or the Image Xpress Micro Confocal high-content imaging system (Molecular Devices).

### Mitochondrial mass and morphology

For mitochondrial mass and morphology, staining with GRP75, TOMM20, and dsDNA was performed (see antibody section for details). 3D image stacks were acquired at optimal Z-distance and reconstructed as Maximum Z-projections. Images were acquired with the Image Xpress Micro Confocal high-content imaging system (Molecular Devices) with a 40x/1.2 N.A. water objective. A TopHat filter was applied to all the channels. The images were thresholded and transformed into a binary segmentation. This segmented area was used to determine mitochondrial morphology parameters.

### Membrane potential determination

Cells were incubated at 37°C (5% CO_2_) for 1 h with 200 nM MitoTracker Green FM (MTG; Thermo Fisher Scientific, M7514), 15 nM TMRE (Invitrogen, T669), and 1 μg/ml Hoechst (Thermo Fisher Scientific, H3570), washed twice with PBS and imaged in phenol red-free media. Images were acquired with the 50 mm slit confocal mode and a 40x (1.2 NA) water lens in Z-stack mode of 0.5 μm slices with a total of 6 slices. Analysis was performed in the MetaXpress software keeping the same parameters for all the images acquired. Maximum Z-projections of MTG were used for morphologic analysis and the sum of Z-projections of TMRE was used for quantification of intensity. A TopHat filter was applied to the MTG images for better definition of structures and equalization of fluorescence. The images were thresholded and transformed into a binary segmentation. This segmented area was used to measure the average intensity of TMRE and the area of mitochondria on the MTG channel. Mitochondria <1 μm^2^ were filtered to count as small mitochondria. The background intensity of TMRE channel was used to set up cut-off intensity value for the depolarized mitochondria.

### Mito-QC imaging and analysis

For mito-QC imaging, images were collected either with the Zeiss LSM880 Confocal system with Airyscan equipped with a Zeiss Pla-Apochromat 40x/1.2 objective, or the Image Xpress Micro Confocal high-content imaging system (Molecular Devices) with a 40x/1.2 N.A. water objective. 3D image stacks were acquired at optimal Z-distance and reconstructed as Maximum Z-projections. A cell-by-cell analysis was performed to count the red vesicles only (mitolysosomes) as described [58]. Briefly, under steady-state conditions, the mitochondrial network fluoresces both red and green; however, upon mitophagy, mitochondria are delivered to lysosomes where mCherry fluorescence remains stable, but GFP fluorescence becomes quenched by the acidic microenvironment. This results in the appearance of punctate mCherry-only foci that can be easily quantified as an index of cellular mitophagy. Images acquired with Airyscan were analyzed using pixel classification segmentation with the Aivia v.10 software. Two segmentations were performed by teaching the pixel classifier to recognize (1) the merged GFP and mCherry signal and (2) the single mCherry puncta signal. Segmentation of mitochondrial network (GFP+mCherry) was used to measure mitochondrial mass while mCherry only pucta was counted as mitolysosomes per cell. For images acquired with the ImageXpress system, a cell-by-cell segmentation was performed using a Gaussian Blur filter of the GFP channel for detection of cytoplasm combined with DAPI detection of nuclei. Cells touching the edges of the field of view (not fully depicted) were excluded from analysis. A range of fluorescence intensity filters were used based on the GFP signal, to exclude high and low signal mitoQC cells; this included the same medium intensity profile cells consistently in the analysis. Following cell selection, a TopHat filter was applied on both GFP and mCherry channels to better define the structures. The images were thresholded and transformed into a binary segmentation, followed by a watershed separation of the mCherry binary segmentation. The binary masks were compared and the mCherry puncta not touching the GFP mask, were counted as mitolysosomes. An average of mCherry pucta per cell was used in all the analyzed images.

### Lysosomal activity and LAMP1 and LC3 distribution

For colocalization analysis, immunofluorescence combining LC3 and GRP-75 or LAMP1 and TOMM20 was performed (see antibody section for details). 3D image stacks were acquired at optimal Z-distance and reconstructed as maximum Z-projections. Images were acquired with the Image Xpress Micro Confocal high-content imaging system (Molecular Devices) with a 40x/1.2 N.A. water objective. A TopHat filter was applied on all the channels. The images were thresholded and transformed into a binary segmentation. This segmented area was used to count LC3 or LAMP1 vesicles, and proximity was determined by overlapping GRP-75 or TOMM20 vesicles, respectively. For lysosomal distribution, the same samples were used to acquire high-resolution images using the Zeiss LSM880 Confocal system with Airyscan. Pixel classification segmentation with the Aivia v.10 software was performed. The distance of lysosomes to the nuclei was calculated to determine the peripheric population of lysosomes defined by the 75^th^ quartile of the mitochondria segmented area to the nuclei.

For lysosomal activity, 10 μg/mL of DQ-Red BSA (Invitrogen, D12051) was incubated for 6 h in the medium at 37°C (5% CO_2_). Under normal conditions, DQ-Red BSA traffics to lysosomes and is cleaved by lysosomal hydrolases, resulting in a bright red fluorescent signal [59]. Disruption of cargo delivery to lysosomes was inhibited with 100 nM Bafilomycin A1. Cells were fixed and stained with LAMP1 for lysosome staining (see antibody section for details). Images were collected with the Image Xpress Micro Confocal high-content imaging system (Molecular Devices) with a 40x/1.2 N.A. water objective. 3D image stacks were acquired at optimal Z-distance. Maximum Z-projections of LAMP1 stained were used for morphologic analysis and the sum of Z-projections of DQ-Red BSA was used for quantification of intensity. A TopHat filter was applied to the LAMP1 images for better definition of structures and equalization of fluorescence. The images were thresholded and transformed into a binary segmentation. This segmented area was used to measure the average intensity of DQ-Red BSA and the area and number of lysosomes on the LAMP1 channel.

### STED microscopy

Live cells were stained with 250 nM Abberior LIVE ORANGE mito dye (Abberior, LVORANGE-0146-2NMOL) for 1 h in phenol red-free medium at 37°C (5% CO_2_), followed by a 20 min wash with medium only. Cells were imaged using the Abberior STEDycon System coupled with a Nikon 100x/1.45 N.A. and STED depletion laser 775 nm. STED images were taken with a 30 nm pixel size and 10 μs dwell time. All the imaging parameters were kept the same between conditions. Analysis was performed by drawing a random line in the middle of the mitochondria and counting peaks of fluorescence to determine the number of cristae per length of mitochondria imaged.

### Time Course Imaging

Following treatment, cells were maintained in an incubator at 37°C (5% CO_2_). Over 48 h, an image was acquired every 4 h by the CellCyte 1. At the completion of the experiment, the CellCyte software was used to create a cell mask to determine the cell confluency. The percentage of confluency was used in downstream analysis.

### RNA sequencing

#### Sample preparation, sequencing, and alignment

Cells from a 10 cm^2^ dish were treated with CAP-1902 for 24 h. After incubation, cells were harvested and pelleted by centrifugation at 300 x g at 4°C for 10 min. Using RNeasy Mini (Qiagen, 74104) and adhering to the manufacturer’s recommendations, RNA was extracted from the pellet and stored at −80°C. Frozen RNA samples were sent to Vanderbilt University Medical Center (Nashville, TN) for analysis. A Quality Control analysis was performed on the RNA samples using the Agilent Bioanalyzer to assess quality. Poly(A) RNA enrichment was conducted using NEBNext Poly(A) mRNA Magnetic Isolation Module (New England Biolabs, E7490), and the sequencing library was constructed by using the NEBNext Ultra II RNA Library Prep Kit (New England Biolabs, E7765L) following the manufacturer’s instructions. End repair, A-tailing, and adapter ligation were performed to generate the final cDNA library. The library quality was assessed using a Bioanalyzer and quantified using a qPCR-based method with the KAPA Library Quantification Kit (KAPA Biosystems, KK4873) and the QuantStudio 12K instrument.

Prepared libraries were pooled in equimolar ratios, and the resulting pool was subjected to cluster generation using the NovaSeq 6000 System, following the manufacturer’s protocols. 150 bp paired-end sequencing was performed on the NovaSeq 6000 platform targeting 50 M reads per sample. Raw sequencing data (FASTQ files) obtained from the NovaSeq 6000 was subjected to quality control analysis, including read quality assessment. Real Time Analysis Software (RTA) and NovaSeq Control Software (NCS) (1.8.0; Illumina) were used for base calling. MultiQC (v1.7; Illumina) was used for data quality assessments.

#### Data analysis

Aligned datasets were filtered and normalized in R using the EdgeR package for differential gene expression contrasts [60]. Gene set Enrichment analysis was carried out with the Fgsea package [61], the REACTOME pathway database [62], the ReactomePA package [63], and the Molecular Signatures Database release 2022.1 [64]. RNAseq-derived figures were generated using the ggplot2 package in R.

### Antibodies

The primary antibodies used were: anti-human OXPHOS cocktail (Abcam, ab11041); anti-PROHIBITIN (Abcam, ab28172); anti-CITRATE SYNTHASE (Abcam, ab96600); anti-MIC60 (Abcam, ab137057); anti-TOMM20 (Sigma-Aldrich, WH0009804M1); anti-phospho AMPK (T172) (Cell Signaling, 2535); anti-AMPK (Cell Signaling, 2532); anti-phospho FUNDC1 (S17) (Invitrogen, PA5-114576); anti-FUNDC1 (Aviva Systems Biology, ARP53280_P050); anti-phospho ULK1 (S638) (Cell Signaling, 14205); anti-ULK1 (Cell Signaling, 8054); anti-LAMP1 (Cell Signaling, 51774); anti-MasR (Santa Cruz Biotechnology, sc-390453); anti-GST-Tag (Cell Signaling, 2624); anti-LC3B (Cell Signaling, 3868); anti-CHOP (Cell Signaling, 2895); anti-LAMIN A/C (Santa Cruz Bio, sc-20681); anti-β3 TUBULIN (Cell Signaling, 5568); anti-VINCULIN (Sigma-Aldrich, V9131); anti-ACTIN (Sigma-Aldrich, MAB1501), anti-TOMM20 Alexa Fluor 488 (Abcam, ab205486); anti-GRP75 (NeuroMab, 75-127); anti-ATF4 (Cell Signaling, 11815); anti-PGC1α (Invitrogen, PA5-72948); anti-SDHA (Invitrogen, 459200); anti-phospho ACC (S79) (Abcam, ab68191); anti-ACC (Abcam, ab45174). For western blotting, primary antibodies were used at 1:1000 except for actin and vinculin, which were used at 1:5000. Primary antibodies in immunofluorescence were used at 1:300 except for LC3B, which was used at 1:100.

The secondary antibodies used were: HRP-linked anti-mouse IgG (Cell Signaling, 7076); HRP-linked anti-rabbit IgG (Cell Signaling, 7074); donkey anti-mouse IgG 488 (Invitrogen, A21202); donkey anti-rabbit IgG 488 (Invitrogen, A21206); donkey anti-mouse 568 (Invitrogen, A10037); rabbit anti-mouse IgG 647 (Invitrogen, A21239). All secondary antibodies were used at 1:3000 for western blotting and 1:300 for immunofluorescence.

### Mitochondria and Crude Organelle Isolation

Primary fibroblasts from confluent 150 cm^2^ dishes were washed and harvested by trypsinization. The pellet was washed twice. Mitochondria were isolated using the Mitochondria Isolation Kit for Cultured Cells (Thermo Fisher Scientific, 89874) according to the manufactureŕs instructions. Briefly, the pellet was resuspended and incubated for 2 min in 400 μL buffer A, supplemented with protease and phosphatase inhibitors, and placed in a glass–Teflon Dounce homogenizer. Cells were homogenized with 80 strokes, the lysate was mixed with 400 μL buffer C and centrifuged at 700 x g for 10 min. The supernatant was transferred to a clean tube and centrifuged at 3000 x g for 15 min for mitochondria isolation and at 20,000 x g for 15 min for crude organelle isolation. The pellet containing the mitochondria or organelles and the respective supernatant fraction were transferred to a clean tube. The mitochondrial pellet was finally resuspended in 50 μl of 1:1 Buffer A:C and the crude organelle pellet in 40 μl of RIPA. Protein concentration was measured by BCA and the fractions were stored at −80°C until further use.

### Nuclear and cytoplasmic fractions preparation

Primary fibroblasts from confluent 150 cm^2^ dishes were washed and harvested by trypsinization. The pellet was washed twice. Nuclear and cytosolic fractions were prepared using the Nuclear Extraction Kit (Novus Biologicals, NBP2-29447) with minor modifications. Briefly, the pellet was resuspended in 0.5 mL of ice-cold 1X hypotonic buffer and transferred to an Eppendorf tube. Cells were incubated on ice for 15 min. 25 μL of 10% detergent was added and the sample was vortexed strongly for 10 s. Tubes were centrifuged for 10 min at 14,000 x g. The supernatant (cytosolic fraction) was transferred to a clean tube and kept on ice. The pellet was resuspended in 50 μL Nuclear Lysis Buffer, vortexed vigorously, and incubated for 30 min at 4°C on a rocking platform. This suspension was vortexed for 30 s and centrifuged at 14,000 x g for 10 min. The supernatant (nuclear fraction) was transferred to a clean tube. Both fractions were stored at −80°C until further use.

### Protein gel electrophoresis and immunoblotting

#### SDS-PAGE

A confluent 10 cm^2^ dish per cell line was washed and lysed using RIPA buffer (0.3 M NaCl, 0.1% SDS, 50 mM Tris, pH 7.4, 0.5% deoxycholate, 1 mM Na_3_VO_4_, 10 mM NaF, 10 mM MgCl_2_, and 1% n-dodecyl-β-D-maltoside, with phosphatase (Thermo Fisher Scientific, A32957) and protease (Thermo Fisher Scientific, 78430) inhibitors. 10–20 μg of either RIPA cell lysate, isolated mitochondria, nucleus, or cytosolic fractions was loaded into 4–12% Bis-Tris gels (Thermo Fisher Scientific, NP0321), and gel electrophoresis was performed in a Mini Gel Tank (Thermo Fisher Scientific) under a constant voltage of 120 V.

#### Immunoblotting

Proteins were transferred to a methanol-activated PVDF membrane (Thermo Fisher Scientific, 88520) in a Mini Trans-Blot cell (Bio-Rad) at a constant voltage of 100 V for 75 min on ice. In blots involving cellular fractions, membranes were incubated with Revert 700 Total Protein Stain (LI-COR Biosciences, 926-11015) according to the manufacturer’s instructions. The total protein signal was imaged in a ChemiDoc Imaging System (Bio-Rad). Then, blots were blocked in 3% bovine serum albumin (BSA; Fisher Scientific, BP9703100) in PBS–Tween20 (1 mL/L; Fisher Scientific, BP337) for 1 h and incubated with the primary antibody overnight. Primary antibodies are listed in the above “Antibodies” section. The next day, membranes were washed three times with PBS-T, incubated with the adequate HRP-conjugated secondary antibody, and washed three more times with PBS-T. Images were acquired in a ChemiDoc Imaging System (Bio-Rad), and band densitometry was quantified using Image Lab (Bio-Rad). When targeting a phosphoprotein, all steps were done using Tris-Tween20 (1 mL/L) buffer.

### Respirometry in intact cells

Primary fibroblasts were plated at 6,000 cells/well in a 96-well Seahorse plate 72 h before the assay. Twenty-four hours after plating, standard maintenance media were replaced with a high concentration of glucose or galactose (4.5 g/L) DMEM supplemented with 2 mM glutamine, 1 mM sodium pyruvate, 1% FBS, and antibiotic–antimycotic. The next day, cells were treated with 5 nM CAP-1902 for 24 h. On the day of the assay, the medium was replaced with Seahorse assay medium composed of DMEM pH 7.4 (Sigma-Aldrich, D5030) supplemented with 5mM glucose or 10 mM galactose, 5 mM HEPES, 2 mM glutamine, and 1 mM sodium pyruvate. OCR and ECAR were measured in a SH XF96 analyzer under basal conditions and after injection of 2 μM oligomycin (Sigma-Aldrich, 495455), two sequential additions of 1.5 μM FCCP (Enzo, BML-CM120-0010), followed by 1 μM rotenone (Sigma-Aldrich, R8875) with 2 μM Antimycin A (AA; Sigma-Aldrich, A8674). Respiratory parameters were calculated according to standard protocols [65, 66], and all rates were corrected for non-mitochondrial respiration/background signal by subtracting the OCR insensitive to rotenone plus AA.

### Statistical analysis

Statistical analyses were performed using GraphPad Prism 10.6.1. All the data shown are the mean ± SEM from at least three biological replicates, represented as individual dots in the graphs. Means were compared using either a two-tailed t-test, one-way ANOVA, or two-way ANOVA. Corrections for multiple comparisons were made by Sidak’s or Dunnett’s multiple comparisons when appropriate. Individual points in a graph denote different biological replicates. Differences were considered statistically significant at P < 0.05, and the exact P-value is annotated in the different panels.

## ACKNOWLEDGMENTS

This work was supported by National Institutes of Health grants: R01 CA232056 (OSS), R01 AA026914 (OSS, ML), R01 DK141923 (OSS). Work was also supported by Capacity Bio (OSS, ML) and the Mitochondria and Metabolism Core (OSS, CB).

## DISCLOSURE AND COMPETING INTEREST STATEMENT

MPO, KE, JG, MD, KG and AW were employees of Capacity-Bio when this study was conducted. M.L. consulted and received a technical support contract from Capacity Bio and is a co-founder of Enspire Bio. KR is a cofounder and consultant for Capacity Bio. KG is a cofounder for Capacity Bio. OSS is a cofounder and SAB member of Enspire Bio LLC, Senergy Bio, and Capacity Bio, and when this study was conducted, he has been serving as a consultant or collaborator to LUCA Science, IMEL, Epirium, Johnson & Johnson, Pfizer, and Stealth Biotherapeutics.

## ABBREVIATIONS

ACE: Angiotensin-Converting Enzyme
ACE2: Angiotensin-Converting Enzyme 2
AMPK: Adenosine 5’-Monophosphate (AMP)-Activated Protein Kinase
Ang (1-7): Angiotensin (1-7)
Ang II: Angiotensin II
AA: Antimycin A
AT1R: Angiotensin II Type 1 Receptor
ATF4: Activating Transcription Factor 4
ATP: Adenosine Triphosphate
ATP6: Mitochondrially Encoded ATP Synthase 6
Bafilo: BafilomycinA1
BCS1L: BCS1 Homolog, Ubiquinol-Cytochrome C Reductase Complex Chaperone
CHOP: C/EBP Homologous Protein
Complex I: CI
Complex II: CII
Complex III: CIII
Complex IV: CIV
Complex V: CV
COX2: Cytochrome C Oxidase Subunit 2
Ctrl: Control
ECAR: Extracellular Acidification Rate
ER: Endoplasmic Reticulum
ETC: Electron Transport Chain
FCCP: Carbonyl Cyanide 4-(Trifluoromethoxy) Phenylhydrazone
FIS1: Mitochondrial Fission Protein
FUNDC1: FUN-14 Domain Containing Protein 1
GFP: Green Fluorescent Protein
GPCR: G Protein-Coupled Receptor
GRP75: Mortalin
ISR: Integrated Stress Response
KD: Knockdown
LAMP1: Lysosome-Associated Membrane Protein 1
LC3: Microtubule-Associated Protein 1A/1B Light Chain 3B
LEUP: Leupeptin + Pepstatin A
MASR: MAS G-Protein Coupled Receptor
MIC60: Mitofilin
mtDNA: Mitochondrial Deoxyribonucleic Acid (DNA)
MTG: MitoTracker Green
OCR: Oxygen Consumption Rate
OMM: Outer Mitochondrial Membrane
OXPHOS: Oxidative Phosphorylation
PGC1α: Peroxisome Proliferator-Activated Receptor Gamma Coactivator 1-Alpha
PMD: Primary Mitochondrial Disorder
RAS: Renin-Angiotensin Pathway
RNA-seq: Ribonucleic Acid (RNA) sequencing
SARS-CoV-2: Severe Acute Respiratory Syndrome Coronavirus 2
SDHA: Succinate Dehydrogenase A
SDHB: Succinate Dehydrogenase B
shRNA: Short Hairpin Ribonucleic Acid (RNA)
STED: Stimulated Emission Depletion
TMRE: Tetramethyl Rhodamine Ethyl Ester
TOMM20: Translocase Of Outer Mitochondrial Membrane 20
ULK1: Unc-51 Like Autophagy Activating Kinase 1
UT: Untreated

## Notes

### Summary of Updates

Disclosure, competing interest statement and affiliations were updated.

