## Supplemental Figures for "Activation of the protective arm of renin-angiotensin system enhances mitochondrial turnover improving respiration and decreasing integrated stress response in a human Complex III deficiency model"

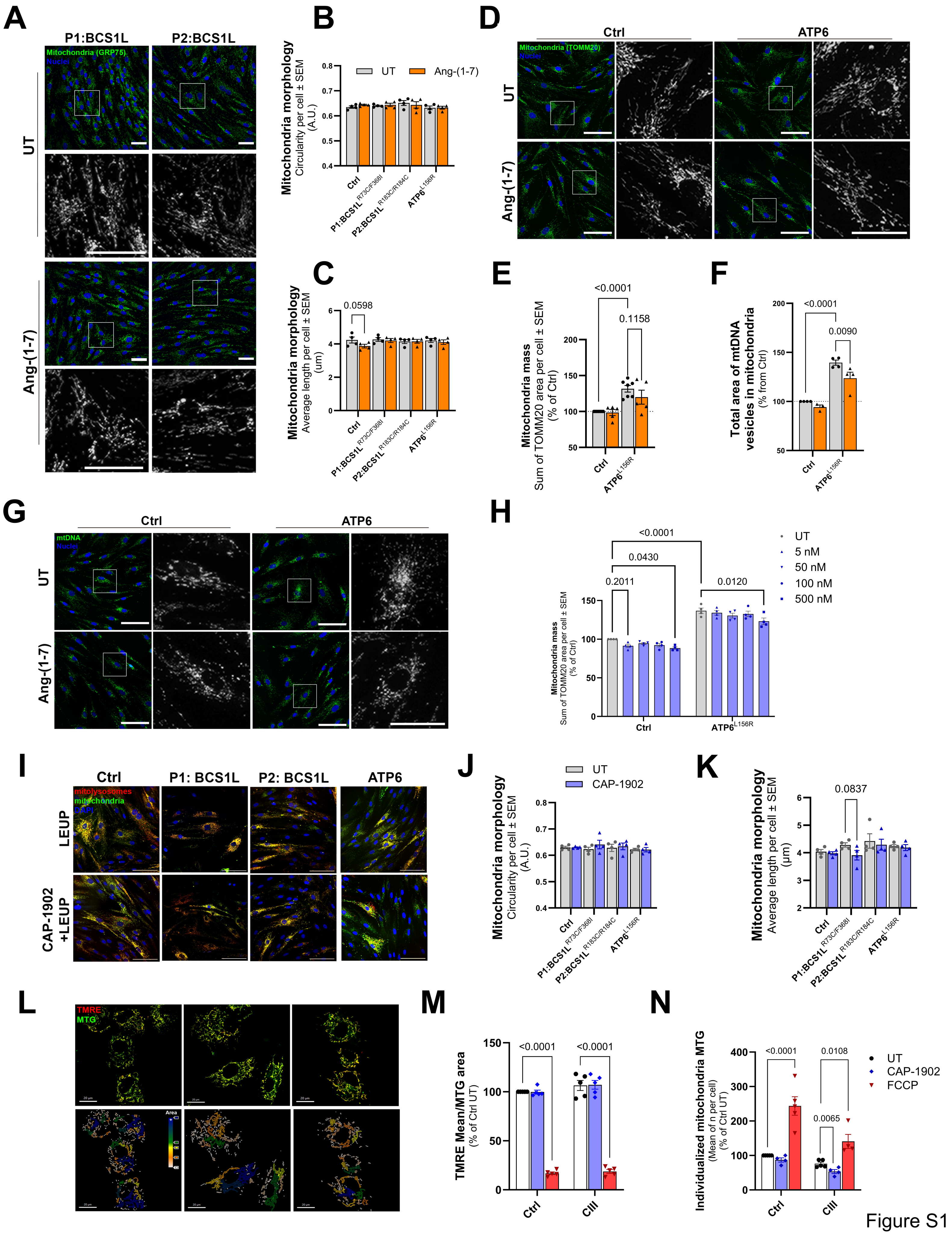
A
P1:BCS1L
P2:BCS1L
Mitochondria (GRP75)
Nuclei
UT
Ang-(1-7)
B
UT
Ang-(1-7)
Mitochondria morphology
Circularity per cell ± SEM
(A.U.)
Ctrl
P1:BCS1L R73C/F368I
P2:BCS1L R183C/R184C
ATP6 L156R
C
Mitochondria morphology
Average length per cell ± SEM
(um)
0.0598
D
Ctrl
ATP6
Mitochondria (TOMM20)
Nuclei
UT
Ang-(1-7)
E
Mitochondria mass
Sum of TOMM20 area per cell ± SEM
(% of Ctrl)
<0.0001
0.1158
F
Total area of mtDNA
vesicles in mitochondria
(% from Ctrl)
<0.0001
0.0090
G
Ctrl
ATP6
mtDNA
Nuclei
UT
Ang-(1-7)
H
UT
5 nM
50 nM
100 nM
500 nM
<0.0001
0.0430
0.2011
0.0120
Mitochondria mass
Sum of TOMM20 area per cell ± SEM
(% of Ctrl)
Ctrl
ATP6 L156R
I
Ctrl
P1: BCS1L
P2: BCS1L
ATP6
mitolysosomes
mitochondria
DAPI
LEUP
CAP-1902
+LEUP
J
UT
CAP-1902
Mitochondria morphology
Circularity per cell ± SEM
(A.U.)
K
0.0837
Mitochondria morphology
Average length per cell ± SEM
(µm)
L
TMRE
MTG
Area
M
TMRE Mean/MTG area
(% of Ctrl UT)
<0.0001
<0.0001
Ctrl
CIII
N
Individualized mitochondria MTG
(Mean of n per cell)
(% of Ctrl UT)
<0.0001
0.0108
0.0065
UT
CAP-1902
FCCP

Figure S1

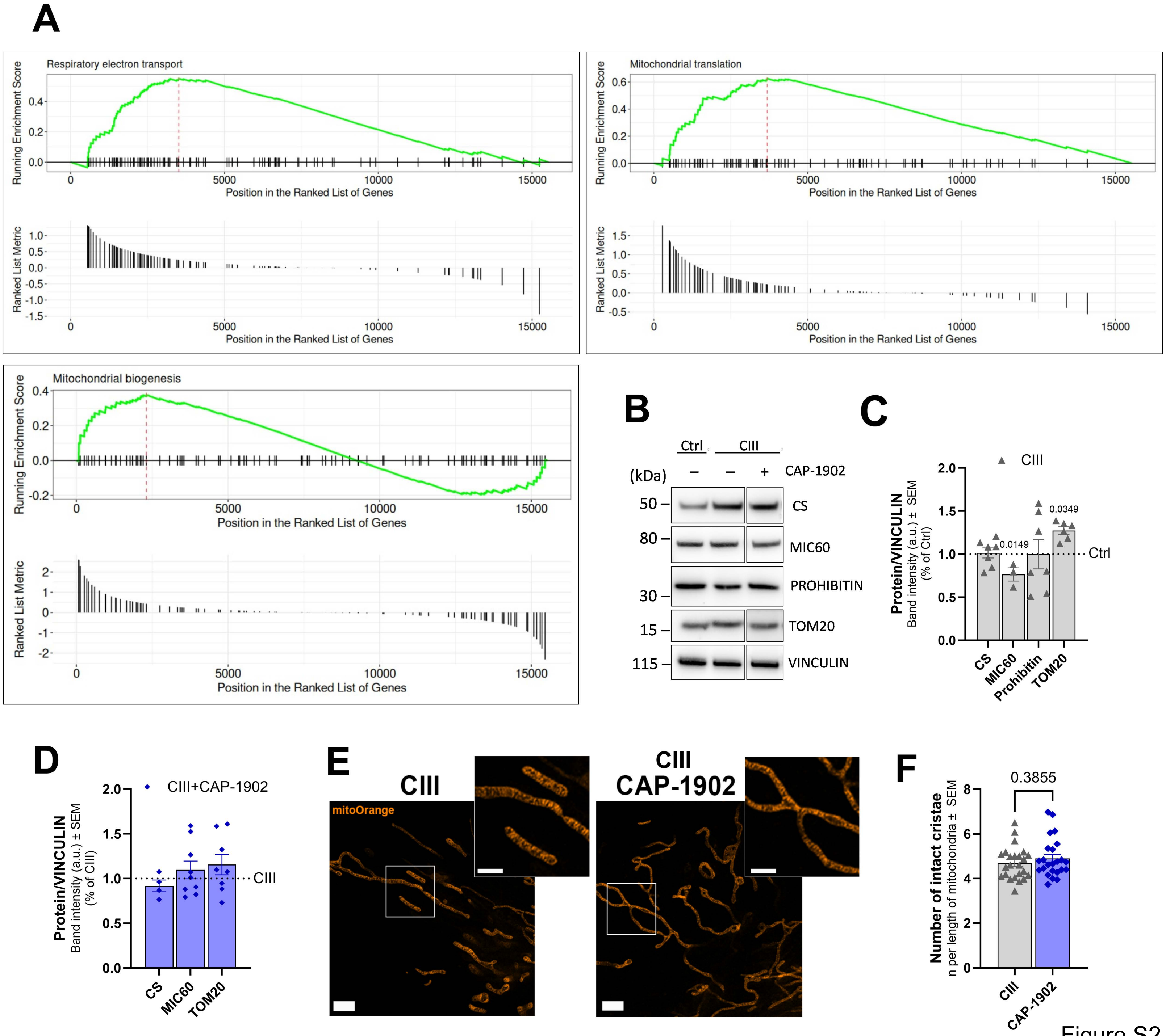
A
Respiratory electron transport
Running Enrichment Score
Position in the Ranked List of Genes
Ranked List Metric
Mitochondrial translation
Mitochondrial biogenesis
B
Ctrl
CIII
(kDa)
CAP-1902
50
80
30
15
115
CS
MIC60
PROHIBITIN
TOM20
VINCULIN
C
CIII
Protein/VINCULIN
Band intensity (a.u.) ± SEM
(% of Ctrl)
0.0149
0.0349
Ctrl
CS
MIC60
Prohibitin
TOM20
D
CIII+CAP-1902
Protein/VINCULIN
Band intensity (a.u.) ± SEM
(% of CIII)
CIII
CS
MIC60
TOM20
E
CIII
mitoOrange
CIII
CAP-1902
F
0.3855
Number of intact cristae
n per length of mitochondria ± SEM
CIII
CAP-1902

Figure S2

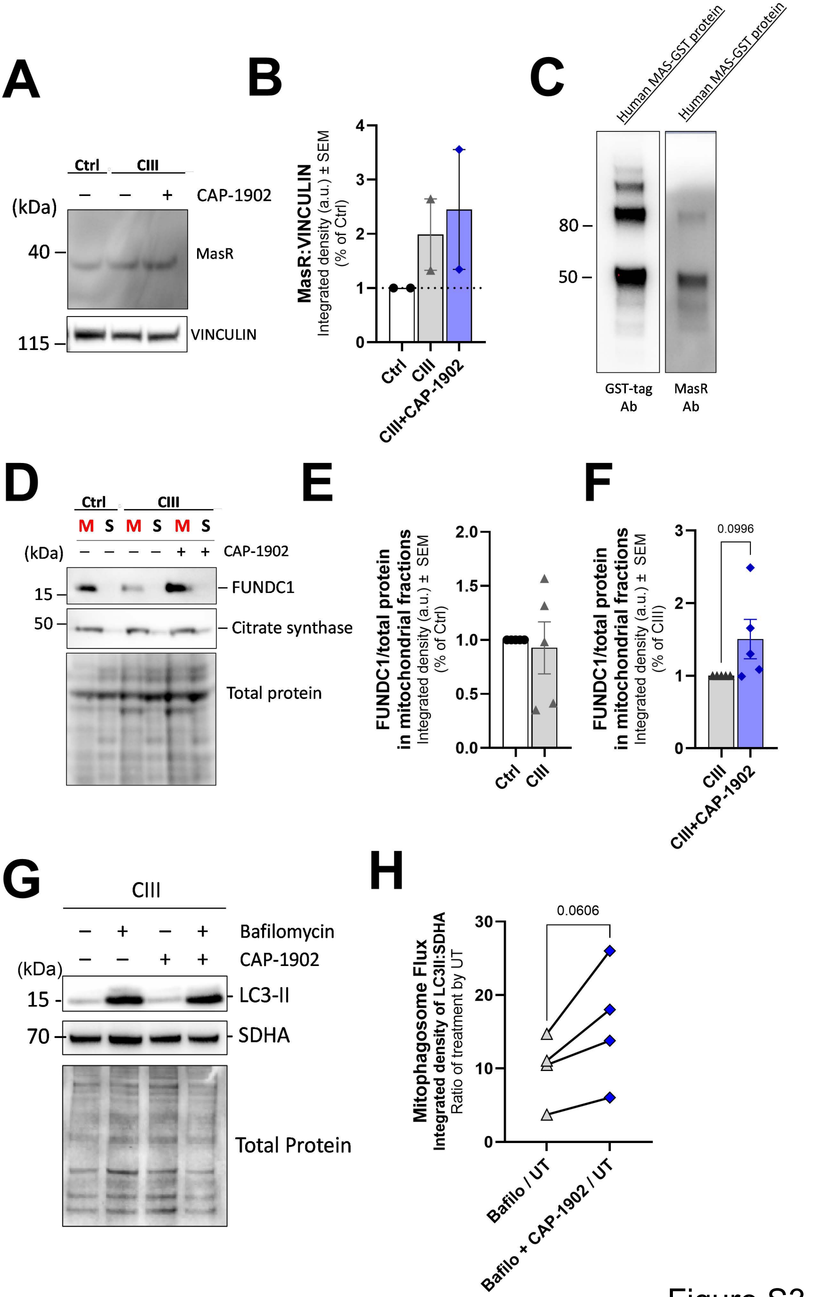
A
Ctrl
CIII
CAP-1902
(kDa)
40
MasR
115
VINCULIN
B
MasR:VINCULIN
Integrated density (a.u.) ± SEM
(% of Ctrl)
Ctrl
CIII
CIII+CAP-1902
C
Human MAS-GST protein
Human MAS-GST protein
80
50
GST-tag Ab
MasR Ab
D
Ctrl
CIII
M S M S M S
(kDa)
CAP-1902
15
FUNDC1
50
Citrate synthase
Total protein
E
FUNDC1/total protein in mitochondrial fractions
Integrated density (a.u.) ± SEM
(% of Ctrl)
Ctrl
CIII
F
0.0996
FUNDC1/total protein in mitochondrial fractions
Integrated density (a.u.) ± SEM
(% of CIII)
CIII
CIII+CAP-1902
G
CIII
Bafilomycin
CAP-1902
(kDa)
15
LC3-II
70
SDHA
Total Protein
H
0.0606
Mitophagosome Flux
Integrated density of LC3II:SDHA
Ratio of treatment by UT
Bafilo / UT
Bafilo + CAP-1902 / UT

Figure S3

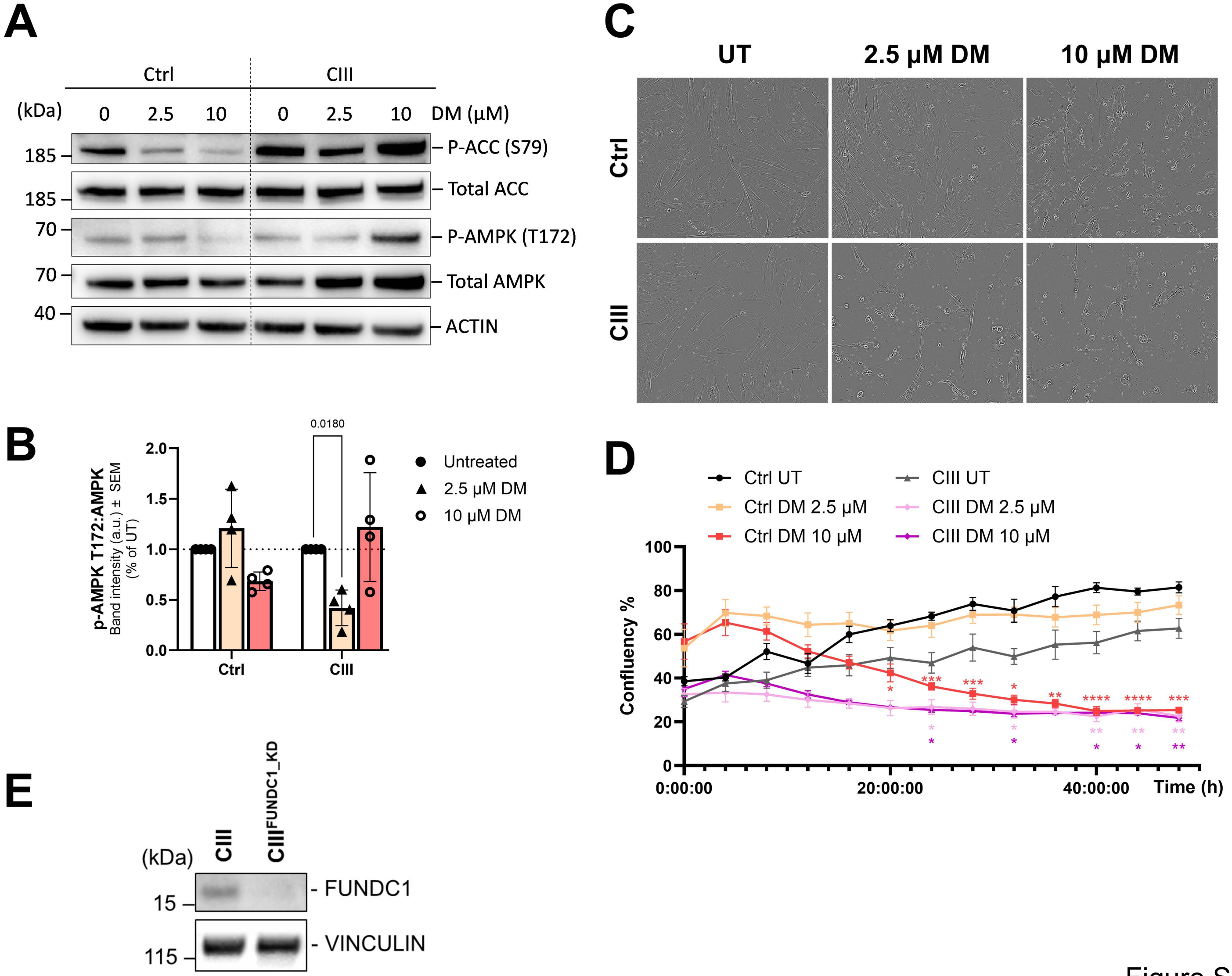
A
Ctrl
CIII
(kDa)
0
2.5
10
DM (µM)
185
185
70
70
40
P-ACC (S79)
Total ACC
P-AMPK (T172)
Total AMPK
ACTIN
B
p-AMPK T172:AMPK
Band intensity (a.u.) ± SEM
(% of UT)
0.0180
Untreated
2.5 µM DM
10 µM DM
C
UT
2.5 µM DM
10 µM DM
D
Ctrl UT
Ctrl DM 2.5 µM
Ctrl DM 10 µM
CIII UT
CIII DM 2.5 µM
CIII DM 10 µM
Confluency %
0:00:00
20:00:00
40:00:00
Time (h)
E
CIII
CIII^FUNDC1_KD
15
115
FUNDC1
VINCULIN

Figure S4

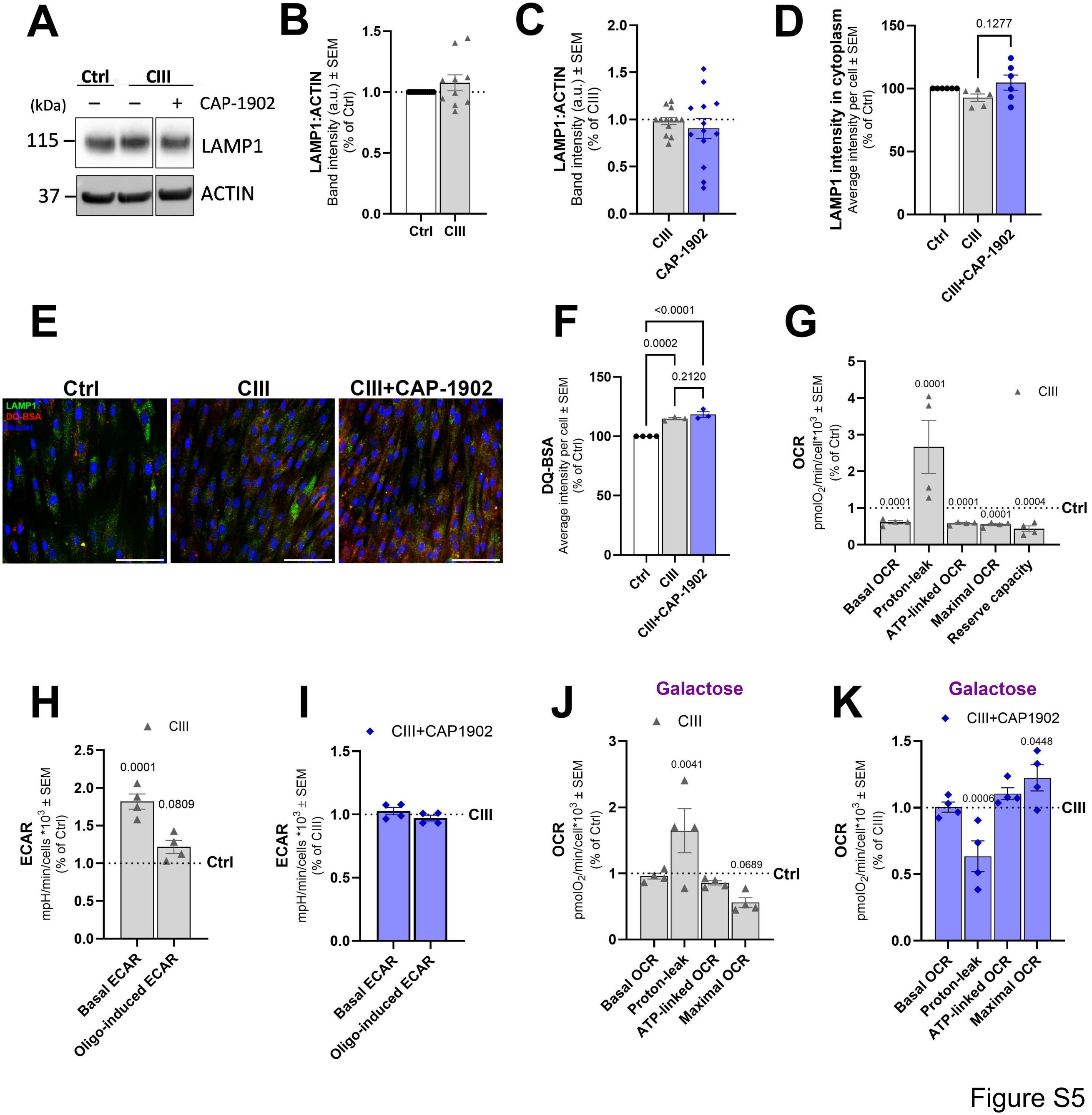
A
Ctrl
CIII
(kDa)
CAP-1902
115
LAMP1
37
ACTIN
B
LAMP1:ACTIN
Band intensity (a.u.) ± SEM
(% of Ctrl)
C
LAMP1:ACTIN
Band intensity (a.u.) ± SEM
(% of CIII)
CAP-1902
D
LAMP1 intensity in cytoplasm
Average intensity per cell ± SEM
(% of Ctrl)
0.1277
CIII+CAP-1902
E
Ctrl
CIII
CIII+CAP-1902
LAMP1
DQ-BSA
Nuclei
F
DQ-BSA
Average intensity per cell ± SEM
(% of Ctrl)
<0.0001
0.0002
0.2120
G
OCR
pmolO2/min/cell*10^3 ± SEM
(% of Ctrl)
Basal OCR
Proton-leak
ATP-linked OCR
Maximal OCR
Reserve capacity
H
ECAR
mpH/min/cells *10^3 ± SEM
(% of Ctrl)
Basal ECAR
Oligo-induced ECAR
I
CIII+CAP1902
(% of CIII)
J
Galactose
K
Galactose
Figure S5

**Figure S1. (A)** Representative high-throughput confocal micrographs of several patient mutant fibroblasts labeled with GRP75 antibody (mitochondria - green), and DAPI (nuclei – blue). Cells were treated with 500 nM Ang-(1-7) for 16 h. Scale bars: 50 μm. **(B-C)** Mitochondrial morphology shown as (**B**) circularity and (**C**) length in patient mutant fibroblast treated with and without Ang-(1-7). Statistical analysis was performed by two-way ANOVA. **(D)** Representative high-throughput confocal micrographs of fibroblasts labeled with TOMM20 antibody (mitochondria - green), and DAPI (nuclei – blue). Cells were treated with 500 nM Ang-(1-7) for 16 h. Scale bars: 50 μm. **(E)** Quantification of mitochondrial mass as the sum of area per cell from TOMM20 images. Statistical analysis was performed by two-way ANOVA. **(F)** Quantification of total mtDNA area as the sum of area per cell from mtDNA images. Statistical analysis was performed by two-way ANOVA. **(G)** Representative high-throughput confocal micrographs of fibroblasts labeled with double stranded-DNA antibody (mtDNA - green), and DAPI (nuclei – blue). Cells were treated with 500 nM Ang-(1-7) for 16 h. Scale bars: 50 μm. **(H)** Quantification of mitochondrial mass as the sum of area per cell from GRP75 images. Ctrl and ATP6 cells were treated for 16 h with CAP-1902 at the concentrations shown. Statistical analysis was performed by two-way ANOVA. **(I)** Representative confocal micrographs of several disease patients expressing mito-QC (mCherry-GFP-FIS1$_{101\text{-}152}$) treated for 16 h with 5 nM CAP-1902 with or without 10 mM Leupeptin/Pepstatin A (+LEUP). Cells were imaged using high-throughput confocal microscopy. Scale bars: 100 μm. **(J-K)** Quantification of mitochondrial mass shown as (**J**) circularity and (**L**) length in mutants with and without CAP-1902. Statistical analysis was performed by two-way ANOVA. (**L**) Representative confocal micrographs of Ctrl, CIII-deficient cells untreated and treated with 5 nM CAP-1902 for 16h, stained with TMRE (red) and MTG (green). MTG was used as a mask (AIVIA segmentation) to create the average TMRE intensity in MTG and select the individualized mitochondria (heatmap for area blue:red; bottom). Scale bars: 20 μm. Cells were imaged with the LSM880 Airyscan microscope in live conditions and analyzed by machine learning pixel classifier in AIVIA software. **(M)** Quantification of averaged intensity of TMRE in the MitoTracker Green area (MTG) in Ctrl and CIII-deficient cells treated with and without 5 nM CAP-1902 for 16 h. 1 uM FCCP was used as a positive control. Statistical analysis was performed by two-way ANOVA. **(N)** Quantification of individualized mitochondria (MTG) in Ctrl and CIII-deficient cells treated with and without 5 nM CAP-1902 for 16 h. 1 uM FCCP was used as a positive control. Statistical analysis was performed by two-way ANOVA. In all cases, data show mean ± SEM. Dots in graphs represent independent biological replicates.

**Figure S2. (A)** Gene Set Enrichment Analysis (GSEA) of mitochondrial pathways in CAP-1902-treated versus untreated CIII cells showing enrichment profiles for three Reactome pathways: Respiratory Electron Transport, Mitochondrial Translation, and Mitochondrial Biogenesis. (**B**) Representative blots showing different mitochondrial markers: Citrate synthase (CS, matrix), MIC60 (IMM), PROHIBITIN (IMM), and TOMM20 (OMM), in Ctrl, CIII-deficient, and CIII-deficient cells treated with 5 nM CAP-1902 for 16 h. Blot images are derived from the same immunoblot and cropped for convenience. (**C**) Quantification of the mitochondria markers in (B) in Ctrl compared to CIII cells. Statistical analysis was performed by unpaired t-test for each protein. **(D)** Average quantification of mitochondrial markers in CIII-deficient compared to CIII-deficient treated with 5 nM CAP-1902 for 16 h. Statistical analysis was performed by unpaired t-test for each protein. In C-D, data shows mean ± SEM of, at least, 3 independent biological replicates. **(E)** Representative confocal micrographs of CIII-deficient cells and CIII-deficient treated for 16 h with 5 nM CAP-1902 and stained with mitoOrange to stain cristae. Scale bars: 2 µm. **(F)** Cells were imaged with the Abberior STEDycon microscope in live conditions and analyzed by tracing a line following the length of the mitochondria and plotting peaks of intensity for counting. Mitochondria cristae were counted and normalized by length of mitochondria. Statistical analysis was performed by unpaired t-test. Dots represent each image analyzed as averaged number of cristae per length of mitochondria from 3 independent experiments. In all cases, data show mean ± SEM. Dots in graphs C and D represent independent biological replicates.

**Figure S3.** **(A)** Representative blot of MasR expression in Ctrl, CIII-deficient, and CIII-deficient cells treated with 5 nM CAP-1902 for 16h. **(B)** Average quantification of MasR protein expression (n=2). (**C**) Antibody quality control confirming the MasR antibody detects the same signal as a GST antibody when detecting a human MasR-GST protein. (**D**) Representative blots of FUNDC1 in mitochondria-enriched fractions (M) vs the corresponding supernatant (S) in Ctrl, CIII-deficient, and CIII-deficient cells treated with 5 nM CAP-1902 for 16 h. Citrate synthase is included as a control for mitochondrial enrichment. **(E)** Quantification of the expression of FUNDC1/total protein in mitochondrial fractions in Ctrl vs CIII-deficient untreated cells and **(F)** CIII-deficient untreated and treated cells with CAP-1902. Statistical analysis was performed by unpaired t-test. (**G**) Representative blots showing LC3-II in crude organelle fractions made from CIII-deficient cells treated with and without 5 nM CAP-1902 and bafilomycin for 16 h. **(H)** Average quantification of LC3II:SDHA representing mitolysosome flux. The effect of bafilomycin in the indicated conditions is highlighted by representing the ratio of the specified treatment by untreated. In all cases, data show mean ± SEM. Dots in graphs represent independent biological replicates.

**Figure S4.** **(A)** Representative blots of phosphorylated (S79) ACC, total ACC, phosphorylated (T172) AMPK, and total AMPK in Ctrl cells treated with 2.5 or 10 μM Dorsomorphin for 30 min. Actin is shown as loading control. **(B)** Average expression of p-AMPK by total AMPK levels in Ctrl and CIII-deficient treated with and without Dorsomorphin. Statistical analysis was performed by two-way ANOVA. Data show mean ± SEM of 3 independent biological replicates. **(C)** Representative images of Ctrl and CIII-deficient cells following 24 h treatment with 2.5 or 10 μM Dorsomorphin. Images were acquired using the Echo Discovery CellCyte. **(D)** Representative cell confluency (in %) of the conditions described in C. Confluency was calculated using CellCyte detection of enhanced contour overtime. **(E)** Representative blot showing the silencing of FUNDC1 in CIII-deficient FUNDC1-knockdown cells. Vinculin was used as loading control. In all cases, data show mean ± SEM. Dots in graphs represent independent biological replicates.

**Figure S5.** (**A**) Representative blots showing the expression of LAMP1 in Ctrl, CIII-deficient, and CIII-deficient cells treated with 5 nM CAP-1902 for 16 h. Actin is used as loading control. Images belong to the same blot but were cropped for convenience. **(B)** Average LAMP1 expression in Ctrl compared to CIII-deficient cells and **(C)** CIII-deficient cells untreated and treated with 5 nM CAP-1902 for 16 h. Statistical analysis was performed by unpaired t-test. (**D**) Average intensity of LAMP1 in cytoplasm obtained from confocal images in Figure S5E. Comparison between Ctrl, CIII-deficient untreated and treated with 5 nM CAP-1902 for 16 h. Statistical analysis was performed by one-way ANOVA. (**E**) Representative high-throughput confocal micrographs of Ctrl, CIII-deficient cells untreated and treated with 5 nM CAP-1902 for 16 h, labelled with LAMP1 antibody (green), DQ-BSA dye (red), and DAPI (nuclei-blue). Scale bar: 200 μm. (**F**) Quantification of DQ-BSA intensity indicating active proteases inside lysosomes (lysosomal activity). Statistical analysis was performed by one-way ANOVA. **(G)** Oxygen consumption parameters in Ctrl compared to CIII-deficient cells, obtained from a Mito Stress Test Seahorse assay. Statistical analysis was performed by two-way ANOVA. (**H**) Average external acidification rates (ECAR) of Ctrl compared to CIII-deficient cells. Statistical analysis was performed by two-way ANOVA**. (I)** ECAR comparing CIII-deficient cells untreated and treated with 5 nM CAP-1902 for 24 h. Statistical analysis was performed by two-way ANOVA**. (J)** Averaged respiratory parameters in galactose conditions of Ctrl compared to CIII-deficient cells. Statistical analysis was performed by two-way ANOVA. **(K)** Respiratory parameters in galactose conditions of CIII-deficient compared to CIII-deficient cells treated with 5 nM CAP-1902 for 24 h. Statistical analysis was performed by two-way ANOVA. In all cases, data show mean ± SEM. Dots in graphs represent independent biological replicates.
